# Covariance-based information processing in reservoir computing systems

**DOI:** 10.1101/2021.04.30.441789

**Authors:** Sofía Lawrie, Rubén Moreno-Bote, Matthieu Gilson

## Abstract

In biological neuronal networks, information representation and processing are achieved through plasticity learning rules that have been empirically characterized as sensitive to second and higher-order statistics in spike trains. However, most models in both computational neuroscience and machine learning aim to convert diverse statistical properties in inputs into first-order statistics in outputs, like in modern deep learning networks. In the context of classification, such schemes have merit for inputs like static images, but they are not well suited to capture the temporal structure in time series. In contrast, the recently developed covariance perceptron uses second-order statistics by mapping input covariances to output covariances in a consistent fashion. Here, we explore the applicability of covariance-based perceptron readouts in reservoir computing networks to classify synthetic multivariate time series structured at different statistical orders (first and second). We show that the second-order framework outperforms or matches the classical mean paradigm in terms of accuracy. Our results highlight a nontrivial relationship between input and reservoir properties in generating the output reservoir activity, which suggests an important role for recurrent connectivity in transforming information representations in biologically inspired architectures. Finally, we solve a speech recognition task for the classification of spoken digits to further demonstrate the potential of covariance-based decoding for real data.

## 1 Introduction

The variability of spiking activity is a hallmark of biological neuronal networks. It has been observed both in vivo and in vitro, in resting and active states and across a wide variety of species, brain areas and time scales (Shadlen and Newsome, 1998; Renart and Machens, 2014; Nogueira et al., 2018). The role that trial-by-trial variability plays in information processing in neuronal networks is currently under debate, after early considerations that saw it as detrimental to (de)coding and learning (Stein et al., 2005; Kostal et al., 2007), but were later proven to not always be the case (Gilson et al., 2011; Boerlin et al., 2013; Moreno-Bote, 2014). In particular, it has recently been shown that structured variability, corresponding to reproducible correlation patterns, can be a substrate for robust information processing (Gilson et al., 2020). That study showed that patterns determined by second-order statistics can be learned and transformed by a simple linear analogue network, thereby implementing a ‘covariance perceptron’.

More generally, covariance coding offers a middle ground alternative between the much-debated rate coding and temporal coding theories (Bair and Koch, 1996; Shadlen and Movshon, 1999; Brette, 2015). In the new view, neither the mean nor the full probability distribution of neuronal activity is used for information transmission, but rather the pairwise correlation between neurons, as quantified by second-order statistics. This view is backed up by recent experimental findings in neurophysiological data, where spike patterns —corresponding to second-or-higher statistical orders— are informative about stimulus or behavior (Panzeri et al., 2017; Shahidi et al., 2019). Considering statistics up to the second order to define informational patterns, we examine how neurons can classify multivariate time series, which has been also used in a variety of applications (Barachant et al., 2013; Sahidullah and Kinnunen, 2016).

In this context, we specifically focus on the processing of patterns determined by structured variability by a reservoir computing system. It usually consists of an untrained network of neurons used to filter incoming signals before feeding a readout layer (Jaeger, 2001; Maass et al., 2002; Lukoševičius and Jaeger, 2009; Tanaka et al., 2019), which draws on the computational power arising from the interplay between recurrent connectivity and neuronal nonlinearities. In particular, nonlinearities can map inputs into spaces where linear separability is easier to achieve, as used in many network architectures like multilayer or deep networks (Hornik, 1991; LeCun et al., 2015). In addition, reservoirs of larger size than the inputs can map them to a higher-dimensional space, which can be beneficial to separate different input patterns. A third point is that, by only training connections from the reservoir to the readout to perform the classification, the weight optimization procedure is simpler (fewer resources to tune) and often more stable as compared to the training of recurrent connections. Reservoir networks thus appear as an interesting candidate to process statistical structures of time series in a classification task, which has been traditionally exploited by reading out the mean activity of the reservoir (Jaeger, 2001; Jaeger et al., 2007; Verstraeten et al., 2007; Lukoševičius and Jaeger, 2009), although other more complex methods have been tried, such as the model space representation (Aswolinskiy et al., 2016).

This study explores the combination of reservoir computing systems with mean/covariance decoding to classify multivariate time series (Fig. 1A, top pathway), using both synthetic and real data. Specifically, we examine the cross-talk between the first and second statistical orders of input times series in the reservoir activity, including both spatial and temporal structure for the second order (i.e. zero-lag and lagged covariances). To do so, we explore through exhaustive numerical simulations and analytically derived insights the influence of the reservoir parameters to elucidate their interplay in shaping the reservoir processing. We implement our reservoirs by means of echo state networks, whose parameters of interest are the spectral radius (quantifying the overall strength of the recurrent connectivity) and the leak rate. These parameters have been shown to influence strongly reservoir properties such as memory capacity for Gaussian inputs (Jaeger, 2002; Farkaš et al., 2016), or the processing of input time series that involve low frequency signals (Verstraeten and Schrauwen, 2009), so we aim to test whether similar tendencies are observed for the processing of second-order statistics.

**Figure 1:**
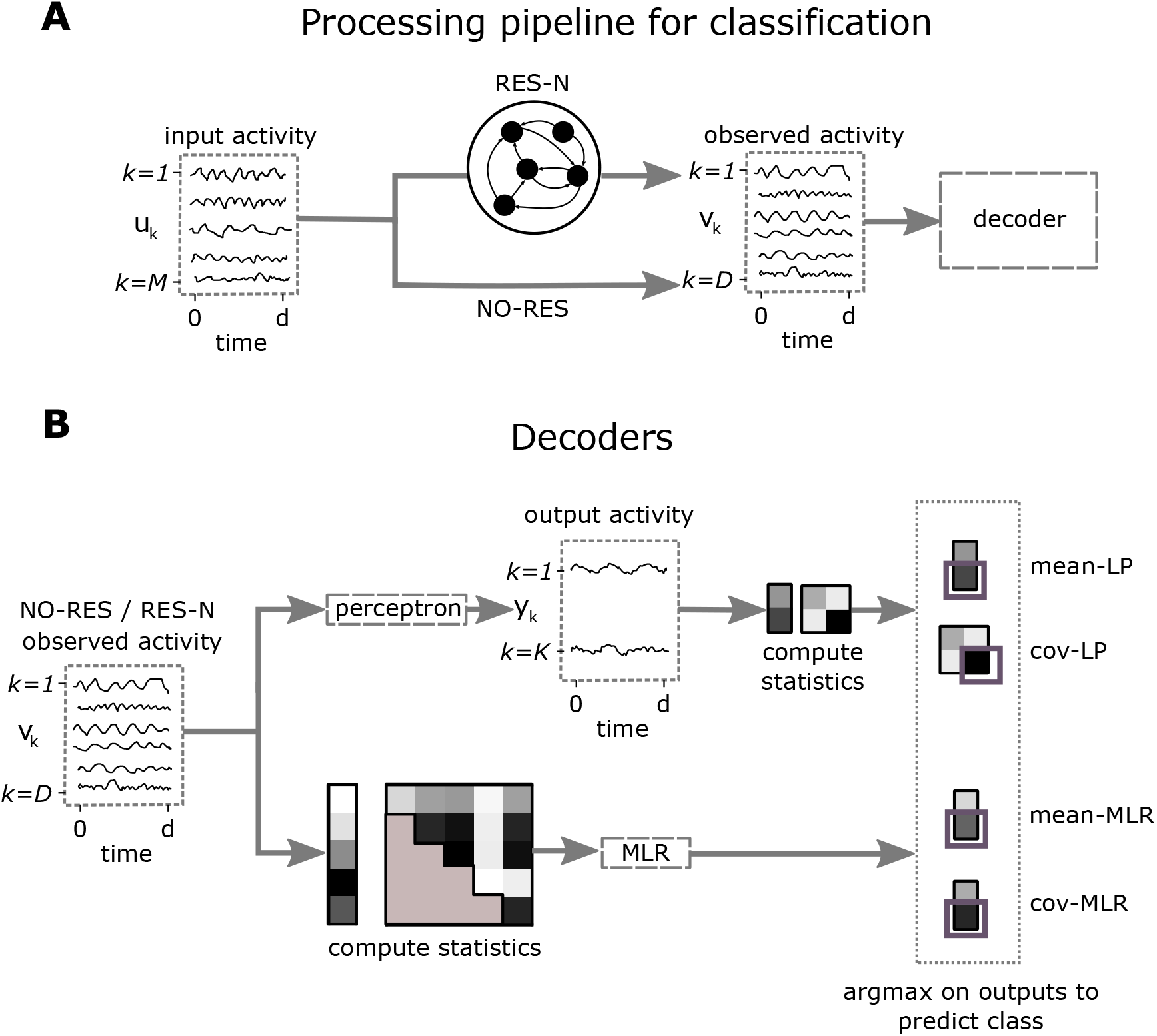
Processing pipeline for multivariate time series classification. **A:** An *M*-dimensional input time series observed for *d* time steps is either filtered by a reservoir layer of *N* neurons (‘RES-N’, top pathway) or directly fed to a linear decoder (‘NO-RES’, bottom pathway) to perform a classification task. The bottom pathway was studied in (Gilson et al., 2020), and we implement it here for comparison purposes. **B:** Our main focus of study is the linear perceptron (LP) decoder (top pathway), but we also implement a multinomial logistic regression (MLR) decoder (bottom pathway) whose accuracy we use as a reference. There is an important operational difference between the LP and the MLR, given by the order in which the statistics are computed for classification. The MLR pathway first computes the statistics of the observed activity, and outputs a class probability vector. This implies the time series are transformed to static features. The perceptron, on the other hand, maps the observed activity time series to output time series, transforming time series to time series. The output time series thus convey information in its statistics that are afterwards evaluated to make the decision

In addition, we compare two types of decoders: a single layer of neurons that are fed from the reservoir inspired by biology (linear perceptron, LP); and the multinomial logistic regression (MLR) coming from machine learning (Bishop, 2006) (Fig 2B). The main difference between these two decoders is that the LP processes its inputs in real time, producing an output time series that has class-dependent structure at the first (mean-LP) or second orders (cov-LP). The MLR, on the other hand, is fed by precomputed input statistics of either first (mean-MLR) or second-order (cov-MLR) and outputs a single class-probabilities vector.

**Figure 2.**
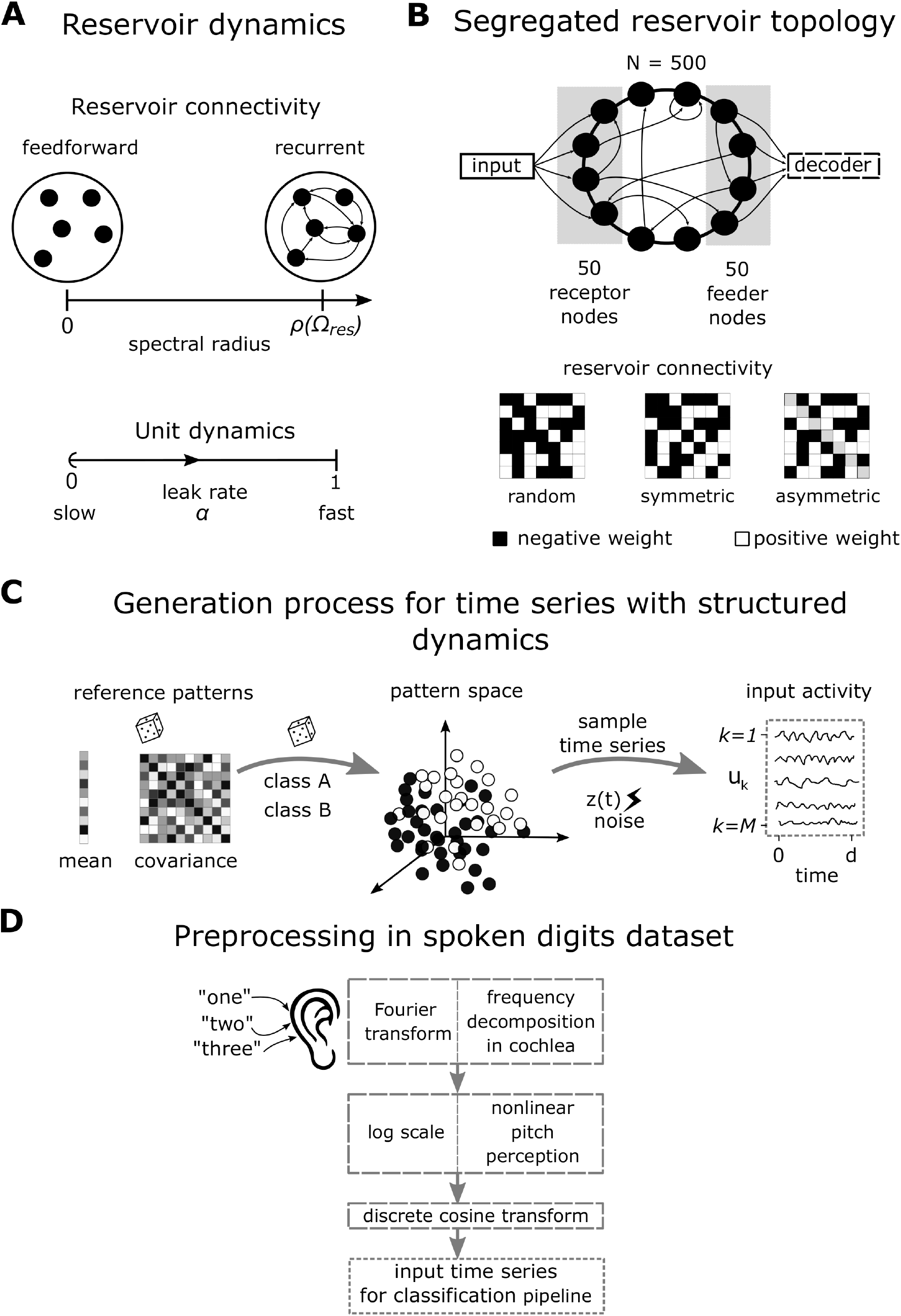
Description of reservoir properties and input datasets. **A:** The reservoir dynamics are characterized by the spectral radius *ρ*(Ω^res^), which is the largest absolute eigenvalue of the reservoir’s connectivity matrix. By convention, a reservoir with *ρ*(Ω^res^) = 0 means Ω^res^ = 0 and corresponds to a feedforward layer without recurrent connectivity. We also modulate the reservoir processing via the local units’ dynamics by adding a leak parameter *α*. Small leak values indicate an integration of the input over time yielding slower update dynamics, while *α* = 1 indicates that the units do not integrate past input information. **B:** We explore an additional reservoir structure where nodes connected to inputs (receptors) are segregated from nodes connected to outputs (feeders) (Kawai et al., 2019). We choose the reservoir connectivity matrix to be fully random, symmetric or asymmetric (see Section 2.1.3). **C:** For synthetic inputs, we first sample reference patterns from given probability distributions. Afterwards, we randomly split them in two balanced classes. To generate a sample time series for a given pattern, we add noise at each time point through specific dynamics (see Eqs. 10-13). **D:** The real data consist of input time series that correspond to spoken digits. To approximate the way the human cochlea processes sound when entering the ear, the speech signals are framed and windowed. For each time bin, a frequency spectrum is computed through a Fourier transform, simulating the frequency-tuning of nerve cells in the cochlea, and a logarithmic scale (mel scale) is used to represent the power coefficients, simulating the nonlinear perception of pitch in humans. These coefficients are decorrelated by means of a discrete cosine transform, keeping 13 amplitude coefficients per time bin. This multivariate time series is afterwards passed through the processing pipeline in B for prediction of the spoken digit. Note that the dataset is only available as preprocessed MFCC coefficients (Hammami and Sellam, 2009; Dua and Graff, 2019).

The manuscript is structured as follows. Section 2 describes the pipeline that we use for classification of multivariate time series, namely the reservoir implementation and the decoders (Fig. 1, followed by the datasets used to test its performance. Results are then presented in Section 3, starting with synthetic time series to uncover general principles and followed by real data to further verify our proposed classification scheme. They are then discussed in Section 4 to contextualize them with respect to their biological and machine-learning implications.

## 2 Methods

This section first presents the reservoir implementation used in our simulations. Among the variety of reservoir implementations that have been proposed (Tanaka et al., 2019), we rely on echo state networks that employ analogue sigmoid neurons (Jaeger, 2001; Jaeger et al., 2007; Verstraeten et al., 2007; Lukoševičius and Jaeger, 2009), since they provide a formalism that is compatible with the covariance perceptron (used as a decoder). We also provide an analysis of the reservoir first and second-order statistics using a weakly nonlinear approximation, inspired by previous work on reservoir dynamics and memory capacity (Aceituno et al., 2020; Verzelli et al., 2020).

Then, we explain the training of the decoder, which is done by performing a gradient descent on its weights (from the reservoir units) in order to minimize the mean-squared error between output activity and target activity as a cost function. The target activity is defined such that the classification can be performed by comparing the values (means or variances) of the readout outputs, in a winner-take-all fashion. Importantly, the gradient descent depends on the metric applied to the reservoir activity and the type of target activity, which differ across the decoders as illustrated in Fig. 1B.

Last, we detail the generation of the input time series that are used to test the classification pipeline. The classification task consists of separating the time series into *K* classes, with *K* = 2 for the synthetic datasets and *K* = 10 for the real dataset. Each of these classes consist of samples that differ by their statistical features, which are transformed by the reservoir and must be captured by the decoder.

We remark that in all the equations to be presented throughout this work, we use lower case greek letters for real parameters, lower case latin letters for vectors and upper case letters (greek or latin) for matrices, except when noted (e.g *M* and *N* are natural numbers).

### 2.1 Reservoir implementation

The reservoir used in the classification pipeline (‘RES-N’ in Fig. 1A, top pathway) is an echo state network with *N* leaky integrator neurons (or units), similar to previous studies (Jaeger et al., 2007; Lukoševičius and Jaeger, 2009). The update equations for the activity state at time *t* of the *N* leaky neurons inside the reservoir, denoted by *x*(*t*) ∈ ℝ^N^ when fed by a multivariate input *u*(*t*) ∈ ℝ^M^, are

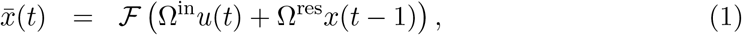

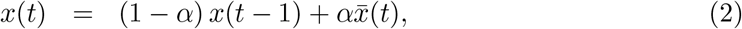

where the function ℱ = tanh has a sigmoidal profile, Ω^in^ ∈ ℝ^N×M^ is the connection matrix from the input time series to the reservoir units and Ω^res^ ∈ ℝ^N×N^ is the weight matrix of recurrent connections within the reservoir. All connection weights are randomly sampled from [−0.5, 0.5), and the resulting matrices are dense. The parameter *α* is a leak rate, *α* ∈ (0, 1], which governs how each reservoir unit integrates its own dynamical state over time. When *α* tends to zero, the neuron’s dynamics becomes slower and more dependent on previous history than on the current input state (Jaeger et al., 2007; Lukoševičius and Jaeger, 2009). When *α* = 1, the activity of the reservoir (Eq. 2) is 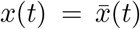, thus, no integration is performed and each unit’s activity state only depends on the instantaneous inputs and activities of other neurons.

Using numerical simulation, we explore the different dynamical regimes of the reservoir, by varying a local parameter (leak rate applied homogeneously to all units) or a global parameter (spectral radius), see Fig. 2A. The spectral radius *ρ* (Ω^res^) is the largest absolute eigenvalue of the reservoir’s weight matrix Ω^res^. It affects the reservoir performance in different benchmark tasks typically reported in the literature, such as memory (Jaeger, 2001; Hermans and Schrauwen, 2010; Farkaš et al., 2016). A general heuristic when *α* = 1 is that *ρ* (Ω^res^) should approximate 1 (from below) for tasks that require long memory and be smaller for tasks where a too long memory might be detrimental (Lukoševičius and Jaeger, 2009). Since we are agnostic to the effects of this parameter for multivariate time series classification (especially those with structured dynamics as we have detailed in the previous section), we vary it spanning a range that goes from 0 to 1.8. In the absence of inputs, a spectral radius larger than 1 implies that a linear reservoir (i.e. when ℱ is the identity operator in Eq. 1) is unstable, in the sense that its trajectory will deviate away from the zero fixed-point when started from a non-zero state (Verstraeten and Schrauwen, 2009). However, in practice, the sigmoid function bounds the growth of the trajectory and effectively produces a reservoir that is dynamically less excitable. Note that, by convention, a null spectral radius implies a zero connection matrix (Ω^res^ = 0), corresponding to a feedforward layer (left-hand side in Fig. 2A). Thus, *α* = 1 and Ω^res^ = 0 implies a nonlinear and memoryless transformation of the inputs randomly mixed by Ω^in^. When *α <* 1, an effective spectral radius can be calculated for the reservoir, corresponding to the linearization of its dynamics: *α*Ω^res^ + (1 − *α*)ℐ_N×N_, instead of Ω^res^ (Jaeger et al., 2007); here ℐ_N×N_ is the identity matrix.

Other possible parameters for exploration include the input scaling and the choice for the sigmoid function ℱ, which we leave for future work.

All the results we present in this study are averaged across 10 different reservoir configurations, where a configuration is given by specific connection matrices Ω^in^ and Ω^res^, and we always start each reservoir from a zero state. Importantly, our work focuses on transient states, since we are interested in learning and representations in short time scales.

#### 2.1.1 Reservoirs propagate diverse input statistics

Under slightly different conditions than the ones we have stated previously (see Supplementary Material A.1 for full details), it can be proven that the first-order statistics of a neuron *x*_*i*_ inside a reservoir with *α* = 1 and spectral radius *ρ* when fed by a multivariate input time series *u*(*t*) ∈ ℝ^*M*^, with *u*_1_(*t*) a bias unit, is given by:

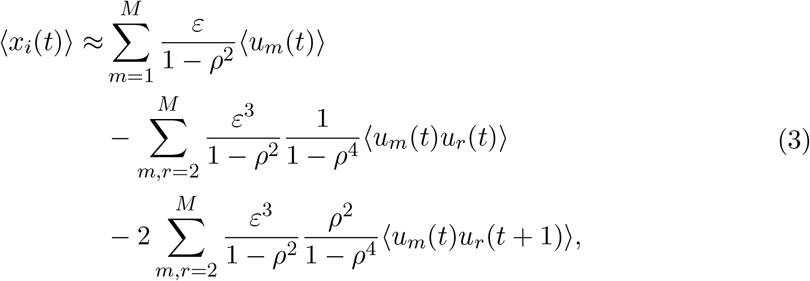

where the angular brackets denote temporal average, *ε* denotes the strength of connections from input to reservoir and we have assumed the neuron is in a weakly nonlinear regime (Supplementary Material A.1).

Likewise, the second-order zero-lag statistics of neurons *x*_*i*_ and *x*_*j*_ in the reservoir are given by:

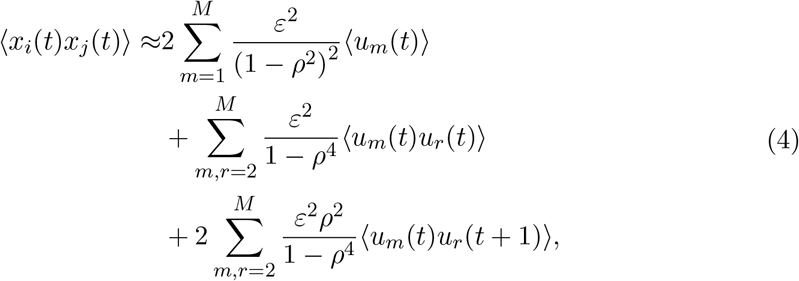

where in the above derivation it is enough to assume that the neuron is behaving in a linear regime (Supplementary Material A.1).

Thus, the reservoir mixes input statistics and these are reflected in reservoir statistics of first and second-order, with a dependence on input-to-reservoir and within-reservoir connections (i.e. *ε* and *ρ*). Full details of this derivation can be found in Supplementary Material A.1. Below we provide some insights for a feedforward reservoir and for the case of temporally correlated inputs.

##### Feedforward reservoir

In the limit *ρ* → 0, the reservoir becomes essentially a feed-forward layer. In this case, reservoir spatial statistics of first and second order do not reflect information embedded in second-order temporal covariances (see Eq. 3 and Eq. 4 when setting *ρ* = 0). Thus, a purely feedforward reservoir is not useful to process this type of structure. Nonetheless, the nonlinear behavior of the sigmoid function introduces a cross-talk between first and zero lag second-order statistics, where we see that the reservoir mean activity and covariances are influenced in both cases by both input means and covariances. Furthermore, in terms of the strength of the input-to-reservoir connections *ε*, first order input statistics have 𝒪(*ε*) contribution to reservoir mean and 𝒪(*ε*^2^) contribution to reservoir covariances, while input covariances have 𝒪(*ε*^2^) contribution to reservoir mean and 𝒪(*ε*^2^). This hints that maintaining consistency betweeen statistical orders for ‘encoding/decoding’ is better than mixed schemes when inputs have first or second-order zero lag structure. Thus, when information about class is embedded in the input mean activity, this is more strongly reflected in the reservoir mean activity but can in principle also be recovered from the (zero-lag) second-order statistics (to a lesser extent). However, when the information is embedded in the zero-lag covariances, the above derivation suggests that second-order statistics of the reservoir provide better representations than first-order ones. This constitutes a strong departure from purely linear networks, where covariance-based information representations are only possible if indeed such a representation is present in the inputs (Gilson et al., 2020).

##### Temporally correlated inputs

This type of inputs require *ρ* ≠ 0. As coefficients of the form 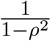 and 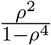 diverge as *ρ* → 1, the optimal value for the spectral radius will be approaching one from below, as it maximizes the influence of input one lag correlation in reservoir mean and spatial covariance statistics. In this limit, from the coefficients in the Taylor series, we observe that reservoir means are more influenced by input lag statistics than reservoir covariances. Note, however, that in this limit the Taylor series approximation is not likely to hold, so we cannot conclude that a mean-based representation be better than a covariance based one.

#### 2.1.2 Analysis of reservoir dynamical regime

Our previous analysis, valid when the reservoir is behaving in a weakly nonlinear regime (see Supplementary Material A.1), points to interesting relationships between reservoir dynamical regimes and useful input-to-reservoir statistics conversions. Therefore, we define for each neuron three possible dynamical regimes according to its activation state *z*, where the activation state is the argument inside the nonlinearity ℱ = tanh at any given time point (Eq. 1). If |*z*| ≤ 0.3, the regime is considered linear. If 0.3 *<* |*z*| ≤ |0.6|, it is weakly nonlinear. Any other case is considered nonlinear. We set these bounds for the regimes by upper-bounding by 0.01 the error in the truncated Taylor series approximation to the hyperbolic tangent of first and third order. For each of the datasets we work with, we numerically compute the probability of finding a neuron in each dynamical regime as function of spectral radius and leak rate.

#### 2.1.3 Reservoir topology

In most of our study we use a reservoir implementation as detailed above, where each neuron is fed by all the input nodes (i.e. Ω^in^ is a full matrix) and likewise feeds the decoder (corresponding to a full Ω^out^ matrix).

However, it is known that the anatomical connectivity in the brain is not full, but sparse and constrained. Getting inspiration from one of the most known pathways of cortex from sensory areas to motor areas, we also consider a reservoir topology that mimics this structure by simply separating input receptor nodes from output feeder ones, as studied in previous work (Fig. 2B) (Kawai et al., 2019). To study how activity propagates across this specialized reservoir, we place the neurons along a ring and choose the input receptor neurons to be opposite the output feeder ones. This implies constrains in Ω^in^ and Ω^out^. For all our simulations using the segregated reservoir, we use a total of *N* = 500 neurons, where two disjoint sets of size 50 form the receptor and feeder groups. Within the reservoir, we try three different naive and sparse (0.1 density) connectivity patterns for Ω^res^: random, symmetric and asymmetric. The motivation behind exploring these different patterns comes from the design of the synthetic inputs in Section 2.3.3, where a mixing matrix given by an asymmetric matrix *J* guarantees, via the exponential function, that the inputs are temporally but not spatially correlated. To generate these topologies, all non-null reservoir connectivity matrix elements are uniformly sampled from [−0.5, 0.5). For the symmetric (asymmetric) matrices, we only sample elements 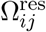 for the upper triangular part, while the lower triangular elements are assigned to fulfill the symmetry (asymmetry) condition 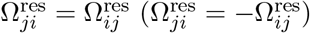.

### 2.2 Decoders

We have four different learning and decoding schemes, as outlined in Fig. 1B: mean-LP, cov-LP, mean-MLR and cov-MLR, with names following the convention ‘feature-classifier’, where the feature corresponds to the statistical order used by the classifier to predict the class. We denote by *v*(*t*) ∈ ℝ^D^ the observed activity in Fig. 1, which either comes directly from the input time series (i.e. *v*(*t*) = *u*(*t*) with *D* = *M*) or is filtered by the reservoir (*v*(*t*) = *x*(*t*) with *D* = *N*). The prefix ‘mean’ indicates that the classifier relies on the mean vector of the observed activity, namely 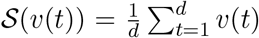. The prefix ‘cov’ indicates that the decoder relies on the matrix of zero-lag (or one-lag) covariances of the observed activity, 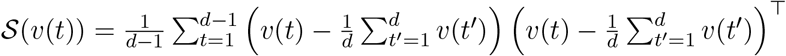, with the superscript ⊤ indicating the matrix transpose. In each case, we consider two options for the classifier: a linear perceptron (Rosenblatt, 1958; Bishop, 2006) (LP in Fig. 1B) or a (multinomial) logistic regression classifier (Bishop, 2006) (MLR in Fig. 1B). Our main focus is the performance of the LP and we implement the MLR decoder only as a reference.

We stress that there is an operational difference between the two classifiers. The LP is biologically inspired in the sense that it generates, for a K-class classification problem, an output time series *y*(*t*) ∈ ℝ^K^ at each time step, and the statistical moments for classification are computed for this vector when the observation period is over (top pathway in Fig. 1B). Thus, the output activity at time *t* is given by:

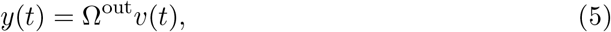

where Ω^out^ ∈ ℝ^K×D^ is the matrix of classifier parameters. We remark that Ω^out^ has a fixed dimension independently of the statistic used for classification. In the mean-based instance, the predicted class is given by the output node with highest mean activity during the observation period. In the covariance-based instance, the predicted class is given by the output node with highest variance during the observation period. Note that when using the covariance perceptron learning rule, we always implement a mapping between spatial covariances, since we do not consider the case of recurrent connections in the readout layer, as is needed to map temporal covariances to spatial covariances (Gilson et al., 2020).

On the other hand, the MLR is a conventional machine-learning approach which first computes the desired time series statistics and afterwards uses it as a vector entry for the classifier (bottom pathway in Fig. 1B). To do so, the MLR produces a single output vector *y* ∈ ℝ^K^ from the statistics 𝒮(*v*(*t*)) of the observed time series, which is not time dependent, but instead represents the probabilities of the input feature vector to belong to each class:

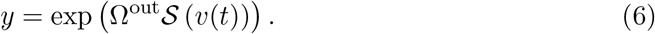

In Eq. 6, 𝒮 (*v*(*t*)) is *D*-dimensional for the mean vector and *D*(*D* + 1)*/*2-dimensional for the vectorized covariance matrix (taking symmetries into account), so the output matrix (Ω^out^) dimension depends on the statistic used as feature. In practice, the output is L1-normalized so that all the elements in *y* sum to 1. The class is then given by the index of the maximum element in *y*. In other words, the order to the transformation by Ω^out^ and the calculation of the statistics is swapped between the two types of classifiers (Fig.1B). Therefore, the MLR for covariances is not equivalent to the linear perceptron with an extra nonlinearity (logistic function). We further emphasize that, once both decoders are trained, while MLR always computes the statistics of the observed activity for classification, the perceptron instead embeds this information directly in the output time series, which are then further processed to compute the statistics and predict the class (Fig. 1B).

In addition, all our models make use of a bias unit at the input-to-reservoir and reservoir-to-output levels, which can be straightforwardly included in all our previous equations. This unit consists of a time series with constant (unit) activity as additional entry. Thus, it has a mean equal to 1, null variance and null cross-covariance with other input nodes. Another possibility would be to choose the bias unit that feeds the decoder unit as a signal with zero-mean activity and variance of 1. Intuitively, this would correspond to adjusting the offset in covariance space. However, to keep consistency at all layers, we keep the bias as a unit constant, given its importance for input representations (see Section 2.1.1 and Appendix A.1).

#### 2.2.1 Learning procedures

Once that the pipeline is set (with or without reservoir, statistical order of feature and decoder), the final step is training the decoder by tuning the matrix weights Ω^out^, the only plastic aspect of the network. In all cases, we rely on a gradient descent that aims to minimize a cost *C*.

For the mean-LP, learning is achieved by minimizing a regularized mean squared cost function between the output mean activity *m* = ⟨*y*(*t*)⟩_*d*_ and a target output mean activity 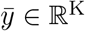 during the observation period *d*:

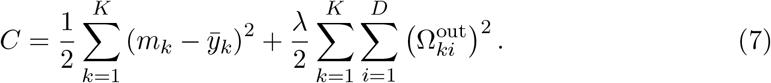

Given the linear nature of the readout, when *λ* = 0, this is equivalent to matching an output time-dependent trajectory to a constant target output trajectory (Gilson et al., 2020), which corresponds to the common winner-take-all readouts typically used in reservoir computing applications for classification (Verstraeten et al., 2007, 2006; Skowronski and Harris, 2007; Jaeger et al., 2007). We use the scikit-learn library (Pedregosa et al., 2011) to minimize this cost function. Importantly, we do not highly tune the regularization parameter *λ*, but set it to 0.02 for all models.

For the cov-LP decoder, learning consists in minimizing a squared error cost function from output spatial (zero-lag) covariances *Y*^0^ ∈ ℝ^K×K^ to target spatial covariances 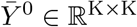 (Gilson et al., 2020):

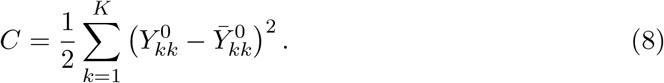

Note that since only the diagonal elements of these matrices are useful for classification, we only constrain them during training, leaving the cross-covariances to vary freely. This is achieved through a gradient-descent learning rule derived for linear dynamics (Gilson et al., 2020), where the weight updates 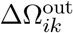 for the connection between element *k* in the output and element *i* in the observed activity are given by:

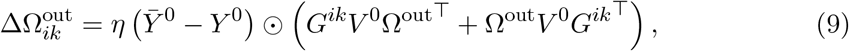

where *η* is the learning rate, *V*^0^ ∈ ℝD×D is the spatial covariance matrix of the observed activity and *G*^*ik*^ ∈ ℝ^K×D^ has 0s everywhere except in element (*i, k*) that is equal to 1. Symbol ⨀ denotes the element-wise (Hadamard) product followed by summation of resulting elements. The learning rate is set for all models to 0.01 and 100 optimization steps are performed.

For the MLR decoders, learning is done through stochastic gradient descent to optimize an L2-regularized cross entropy cost function (Bishop, 2006). For this, we use the scikit-learn library (Pedregosa et al., 2011).

#### 2.2.2 Subsampling procedure for observed activity

Generally, statistical models with larger number of free parameters will yield better performing models than those with lower resources, provided they do not overfit the data (Bishop, 2006). Since the cov-MLR decoder differs in the number of parameters to learn when compared to the other three schemes, to fairly compare them we subsample the dimensions of the observed activity vector *v*^(^*t*) so that its vectorized covariance matrix has dimension close to *D*. In the reservoir pipeline, *D* represents the size *N*. Thus, if *N* = 25, we train the cov-MLR decoders with neuron subsamples of size *S* = 6 and *S* = 7. The resulting decoders have, on average, 24.5 free parameters, thus approximately matching the complexity of cov-LP, mean-LP and mean-MLR when trained on the full size reservoir. Each reservoir initialization is randomly subsampled 100 times for each *S*, so reported performance is not heavily dependent on a given subsampling (as it is averaged across 10 reservoir initializations, with a total of 2, 000 subsampling iterations). For *N* = 50 we use *S* = 9, 10, and for *N* = 100, *S* = 13, 14. For the pipeline without reservoir, we subsample the number of inputs following the same approach. Therefore, for the real dataset with 13 input features (Section 2.4) we use *S* = 4 and *S* = 5.

### 2.3 Synthetic datasets

This section introduces the synthetic datasets of multivariate time series whose ‘information’ relevant for classification is embedded in one of their statistics up to second order. We also consider a mixed scenario where the information is embedded both in means and zero-lag covariances, thus either of these statistics can be used for classification. Note that we use the term ‘information’ in a colloquial manner in this study, without a specific reference to information theory.

We consider a multivariate time series given by *u*(*t*) ∈ ℝ^M^ that represents the activity of *M* = 10 input nodes observed at discrete times 1 ≤ *t* ≤ *d* with *d* = 20. We rely on dynamical systems to enforce a specific spatio-temporal structure that constrains its statistics up to the second order, namely, the empirical mean activity and their zero-lag and one-lag covariance matrices, defined as follows:

- vector of mean activity *p* ∈ ℝ^M^, with elements 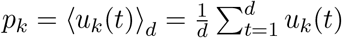
- zero-lag covariance matrix *P*^0^ ∈ ℝ^M×M^, with elements

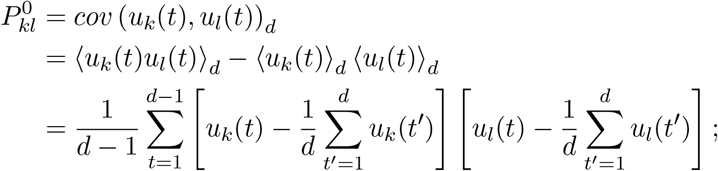
- one-lag covariance matrix *P*^1^ ∈ ℝ^M×M^, with elements

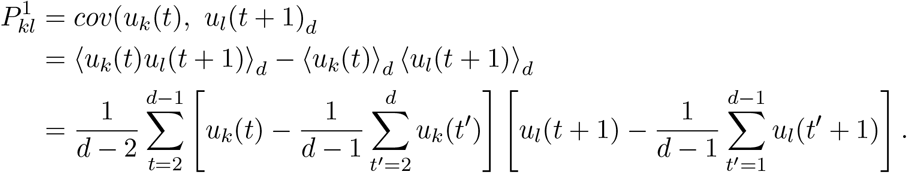

Here our goal is to classify the synthetic time series into *K* = 2 classes. We thus generate two groups of such time series according to one of the above structures, where the defining statistics —*p, P*^0^ and *P*^1^— correspond to distinct patterns that are randomly assigned to one of two classes (Fig. 2C). We first draw a number of such “reference” patterns by sampling given probability distributions, then we generate for each pattern several sample time series for our classification task (each sample involving further stochastic randomness). In each category, there are then two sources of “noise” or variability: the different patterns belonging to a same class, and the empirical noise due to the individual probabilistic realization of each sample. The rationale behind our choice is to account for empirical noise that is typically observed in real time series, such as speech sounds, where distinct phonemes have a similar spatio-temporal structure, which is altered at each pronunciation.

We use cross-validation to assess the classification performance, relying on a 70/30 train/test split that is maintained in all synthetic datasets. For each dataset, we evaluate how separable the two classes are in the relevant feature space by applying a multinomial logistic regression decoder (details in Section 2.2) directly on the input sample time series statistics(see bottom pathways in Fig. 1). The performance of this benchmark decoding is affected by the number of patterns to classify per class (intuitively, densely populated feature spaces are more likely to involve overlapping classes) and by the properties of the probability distribution the patterns are drawn from. Note that we focus on synthetic datasets with a benchmark classification accuracy below 100%, so we can detect performance improvements and decreases across different decoding schemes and models.

In the following subsections we describe the generative dynamics for each statistical structure.

#### 2.3.1 Mean or first-order structure

We use the following generative process for the time series:

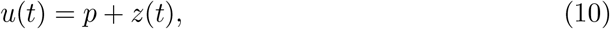

where *z*(*t*) ∈ ℝ^M^ is a normally distributed random variable, with zero-mean and identity covariance matrix. To characterize the mean activity of *u*(*t*), we use a pattern vector *p* ∈ ℝ^M^, with non-null elements sampled from a zero-mean and unit variance normal distribution. This vector is created sparse, with a 0.1 density. This means that 90% of the input nodes will have zero-mean activity.

To generate a dataset of this type, we first draw 20 patterns of such *p* vectors and randomly split them in two classes (10 in each), as shown in Fig. 2C. Once defined the patterns for the two classes, we generate noisy samples by using Eq. 10 with 500 repetitions for each pattern. From each set of 500 samples, we use 350 samples for the training set and the remaining 150 for the test set.

#### 2.3.2 Spatial covariance or second-order zero-lag structure

We use the following generative process for the time series:

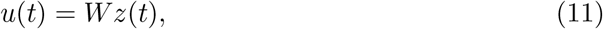

where *z*(*t*) ∈ ℝ^M^ is, as before, a standard normal random variable and *W* ∈ ℝ^M×M^ is a random sparse matrix with 0.1 density and non-zero elements sampled from a standard normal distribution. The resulting zero-lag covariance matrix is given by *P*^0^ = *WW*^*T*^ (Gilson et al., 2020). Note that the generated time series has zero-mean activity over time up to the empirical noise, as well as zero temporal correlations (*P*^1^ = 0). Thus, the only discriminative information for the binary classification is in *P*^0^ (or equivalently, *W*), which is the defining statistic.

To generate a dataset of this type, we sample 60 *W* matrices and randomly split them in two classes before simulating the dynamical processes. As before, we then generate 500 noisy samples with a 70/30 ratio for the train and test.

#### 2.3.3 Temporal covariance or second-order one-lag structure

We use the following generative process for the time series:

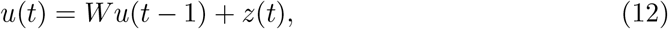

where we choose the mixing matrix *W* = exp(*β*ℐ_M×M_ + *J*), with parameter *β <* 0, ℐ_M×M_ ∈ ℝ^M×M^ is the identity matrix and *J* ∈ ℝ^M×M^ is an antisymmetric matrix. This guarantees that the time series will not differ neither in their mean activity vectors *p* (which are null) nor in their spatial correlation structure *P*^0^ (which only depends on *β*), but only in their one-lag covariances *P*^1^ = *WP*^0^ (Gilson et al., 2020).

To generate a dataset of this type, we sample 6 *W* matrices and randomly split them in two classes before simulating the dynamical processes to generate the noisy samples in the same manner as before. Without loss of generality, we set *β* = −0.5 and create the matrices *J* by sampling unsigned upper diagonal elements from the uniform distribution over [0.5, 1). The elements’ signs are randomly assigned, and the resulting *J* matrices have by construction a 0.3 density.

#### 2.3.4 Mixed spatial inputs with first and second-order zero-lag structure

To create time series that differ in mean and spatial covariance structure, we use a superposition of the signals given in Eqs. 10 and 11:

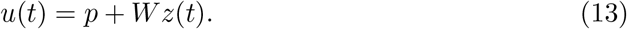

To generate a dataset of this type, we randomly sample 20 *W* matrices and 20 *p* vectors. Each *p* is randomly paired to a *W* matrix and the tuple is randomly assigned to one of two classes. This is done with the purpose of having mean patterns and covariance patterns that are evenly separable. Afterwards, the dynamical processes are simulated. Note that *p* and *W* are generated with the same density and normal distribution for their non-zero elements, as in Sections 2.3.1 and 2.3.2 respectively.

### 2.4 Spoken Arabic digits dataset

To test our covariance-based decoding applied to reservoir computing in a real application, we work with the spoken Arabic digits dataset (Hammami and Sellam, 2009; Hammami and Bedda, 2010; Dua and Graff, 2019). The motivation is to use time series with spatio-temporal structures. The dataset contains 8800 multivariate time series (10 digits x 10 repetitions x 88 speakers) recorded from native Arabic speakers (44 females, 44 males, ages 18-40 years old) with the purpose of classification, split in a training (75%) and test set (25%). Here the classification is not binary, but there are *K* = 10 classes (one per digit).

The time series are represented by 13 Mel Frequency Cepstral Coefficients (MFCC) (Davis and Mermelstein, 1980), which constitute a widely used feature for tasks such as speech recognition (Usman, 2017). They mimic the transformation of the audio signal by the inner ear and are a model of how sound stimuli are “perceived” by the early neuronal auditory system. As represented in Fig. 2D, when a mechanical sound wave reaches the ear, it produces vibrations that propagate throughout the cochlea, with high frequencies entraining the early part of the cochlea and low frequencies the end part. Hair cells in the cochlea translate these vibrations into electrical activity in a frequency dependent manner (depending on their position), so the sound is spectrally decomposed. This is performed by means of the Fourier transform in the MFCC computation. Afterwards, the spectrum is represented using a logarithmic scale (mel scale), emulating the nonlinear perception of pitch. Finally, a discrete cosine transformation is applied with the purpose of decorrelating the resulting coefficients.

Since the MFCC sequences vary in length (5-92 elements, median 40), as is natural in speech, we opt to shrink or expand them (through zero-padding) to have the same size and a consistent observation window (*d* = 50). This is strictly not necessary since all our classification methods can operate on varying sequence lengths, but is only motivated to make the implementations more straightforward and does not make the classification problem easier. Another possibility to avoid losing the information about sequence length, which strongly relates to digit identity, is to add it as a normalized constant input to the reservoir (Jaeger et al., 2007), but we choose to not follow this approach.

## 3 Results

The purpose of our study is the comparison between mean-based and (co)variance-based linear perceptron readouts applied to a reservoir of sigmoid neurons for the classification of multivariate time series. To do so, we firstly consider synthetic time series with controlled structures, characterized by either their means, spatial covariances (zero-lag) or temporal covariances (one-lag) (see Section 2.3).

To determine the usefulness of the reservoir in the pipeline for classification, we also compute baseline classification accuracies (i.e. without reservoir, ‘NO-RES’ pathway in Fig. 2A) for each perceptron readout type. Furthermore, we use additional multinomial logistic regression decoders (mean-MLR and cov-MLR) to quantify in terms of their accuracy how ‘class-informative’ the noisy statistics of the time series that reach the perceptron readouts are. We remark that the purpose of the MLR decoders is merely to provide a reference, and that these decoders are operationally different from the perceptron. The later receives time series as input and likewise produces time series as output, with class ‘information’ embedded in its statistics. MLR instead receives pre-computed statistics as input features and directly outputs a single static class-probability vector (Fig.1B and see Section 2.2 for further details).

We systematically vary reservoir parameters to allow for the identification of which properties of the reservoir (size, spectral radius, leak rate) are important to extract the relevant information for classification, in line with previous work that used mean-based readouts with similar reservoirs (Jaeger, 2001; Farkaš et al., 2016; Boedecker et al., 2012; Schaetti et al., 2016). In addition, we explore the influence of the reservoir connectivity by comparing fully connected random reservoirs and segregated reservoirs where input-receptor neurons (receptors) and output-sending ones (feeders) are separated by at least one neighbor (Fig.2B and see Section 2.1) for the decoding of inputs with different structures.

Last, we apply our analysis under the same considerations of the first part of our study to real data for speech recognition (see Section 2.4), which is a practical problem where reservoir computing has been efficiently applied (Verstraeten et al., 2005, 2006, 2007; Skowronski and Harris, 2007; Jaeger et al., 2007; Triefenbach et al., 2010; Alalshekmubarak and Smith, 2014; Zhang et al., 2015; Jin and Li, 2017).

### 3.1 Reservoirs enhance covariance perceptron performance and efficiently represent second-order statistics from input time series

The linear perceptron readout (mean-LP and cov-LP in Fig. 1B) is trained to perform a binary (*K* = 2) classification task of synthetic input time series that differ by their second-order statistics. We create these time series in such a way that the statistical features that determine to which of the two classes they belong to is embedded either in their zero-lag covariances (spatial structure, Section 2.3.2 in Methods) or in their one-lag covariances (temporal structure, Section 2.3.3 in Methods).

For the spatial structure where the categories differ by the zero-lag covariance patterns, we first train mean-LP and cov-LP readouts without the reservoir in the pre-processing stage. As expected, mean-LP operating directly on the input time series does not produce above chance classification accuracy, since the inputs have, by construction, zero-mean activity (up to some observation noise). On the other hand, cov-LP achieves an accuracy of 71 ± 2%. While this accuracy is much better than chance, the decoder is not able to fully extract the second-order information embedded in the input time series covariances, as quantified by the cov-MLR performance (88.6 ± 0.9%).

We add the reservoir to the pipeline and vary its parameters (namely size, leak rate and spectral radius) to assess how they influence the classification performance (Fig. 3A, left panel). For the classical mean-based perceptron readout (in blue), we find similar results to previous work on reservoir computing applied to others tasks (e.g. types of inputs): performance monotonically increases with the number of units forming the reservoir. This is due to the increased dimensionality of the representation of the inputs in the reservoir activity, which makes it easier to find a separating hyperplane for the two categories of inputs. The new approach with the cov-LP readout (in red) displays the same trend. Furthermore, the reservoir can boost the decoding performance beyond that of a cov-LP directly applied to the inputs (dark gray line) with as little as *N* = 25 neurons (half the amount needed by mean-based readouts) and it even reaches the performance of the MLR directly applied on input covariances (light gray line) for *N* = 100. Note, however, that the number of trained weights per class is then equal to *N* = 100 for the cov-LP, whereas it is equal to 10 × 11*/*2 = 55 for the (cov)MLR-NO-RES (see Section 2.2). For both readout types (mean/cov), we find that the performance decays with spectral radius, the best being achieved when the reservoir is a feedforward layer where *ρ*(Ω^res^) = 0. This points to interesting relationships between reservoir dynamics and representations. Indeed, it can be shown that when the neurons inside the reservoir behave linearly, then mean-based reservoir representations are not able to classify spatially structured inputs at the second-order, while covariance-based ones can (see Section 2.1.1). However, when neurons are driven in a weakly nonlinear regime, mean-based representations become possible. On the other hand, when a reservoir is excited at a strongly nonlinear regime (i.e. saturating the nonlinearity), it will provide representations (at both orders) that are degraded when compared to the weakly nonlinear case.

**Figure 3.**
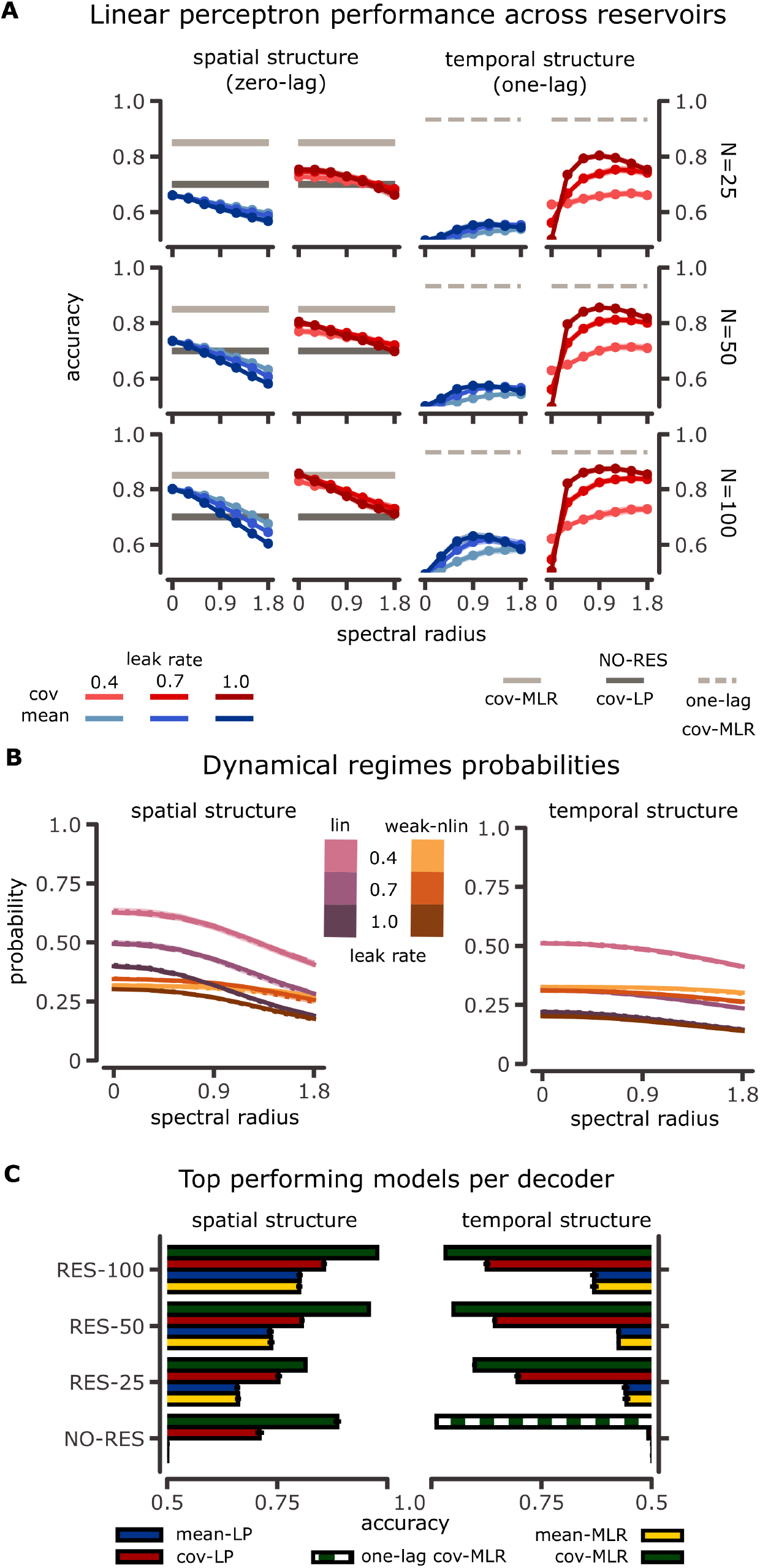
Decoding performance for spatial and temporal structure and reservoir dynamics. **A:** Reservoir classification performance for a spatial structure (left panel) and a temporal structure (right panel) embedded in the input time series, when the decoder is a linear perceptron. Accuracy is shown as a function of spectral radius (on the x-axis) and for different reservoir sizes (see *N*) and leak rates (indicated by the various contrasts). We compare mean-based decoder (mean-LP in blue) and covariance-based decoder (cov-LP in red). The light-gray lines on each subplot indicate the performance of a MLR classifier directly applied to the statistics (see bottom pathways in Fig. 1) that characterizes the information embedded in the input time series (zero-lag covariances for the left panel and one-lag covariances for the right panel). The dark gray line shows the performance of a covariance perceptron directly applied to the input time series (top pathways in Fig. 1). Shaded areas represent ±1 standard error of the mean (sem) across 10 different simulations of time series and reservoir configurations. Each class has 30 different covariance patterns in the left panel, and 6 in the right panel. **B:** Probability of finding a reservoir neuron in the linear dynamics regime (see Section 2.1.2) (pink) or in the weakly nonlinear regime (orange) versus spectral radius, for different reservoir sizes (overlapping dotted, dashed and solid lines for *N* = 25, *N* = 50 and *N* = 100 respectively) and leak rates (shades). Results are averaged across 10 different data simulations and reservoir configurations. Shaded areas are ±1 sem. **C:** Accuracy for the best models for each decoder type: blue for mean-LP, red for cov-LP, yellow for mean-MLR and green for cov-MLR. Error bars represent ±1 sem across 10 different simulations of time series and reservoir configurations. Numbers in model names indicate the size of the reservoir (RES-*N, N* = 25, 50, 100 as above, see the bottom pathway in Fig. 2A), while NO-RES indicates that the input time series is directly fed to the decoder (the bottom pathway in Fig. 1A). In the right panel, we also include the performance of MLR on one-lag covariances (one-lag cov-MLR) for the NO-RES case, since the other features are not informative in this case.

In fact, a neuron saturating the nonlinearity can only behave in three possible ways: continuously saturating the nonlinearity at the top limit, at the bottom one, or alternating from one to the other. Neurons constantly saturating the same limit will have average state equal to 1 or -1, while flipping neurons will have mean states that depend on the switching probability, with higher probabilities more likely leading to zero-mean states that degrade the input representation. This degradation, nonetheless, is expected to be more prominent in the mean than the covariance space, since saturated neurons can display coordinated switching behavior.

Succinctly, when the goal is to process input spatial covariances, randomly mixing inputs (via Ω^in^) and applying a point-wise nonlinearity is enough to achieve this, while keeping a memory of past states through reservoir dynamics appears detrimental. Indeed, strong recurrent connections within the reservoir drive it away from the linear/weakly nonlinear regime (Fig. 3B) and the corresponding accuracy drops as representations lose quality by becoming sparser. In line with this, performance degrades more slowly for the cov-LP than the mean-LP, although the effect becomes similar as reservoir size increases.

We also observe that cov-LP performance is largely insensitive to changes in the leak rate, as shown by the overlapping red lines in Fig.3A (left panel). On the other side, mean-LP performance degrades when increasing the leak rate for reservoirs whose spectral radius is close to or larger than 1, and this effect increases with reservoir size. Thus, the leaky integration mechanism becomes useful when using mean-based readouts (smaller *α* values yield better performance). Indeed, leaky integration drives the reservoir away from the nonlinear regime, therefore improving representations (see profiles in 3B with varying leak rate).

Then, we consider the temporal structure signal (Fig. 3A, right panel, see Section 2.3.3 in Methods for details) for which the input time series from the two classes only differ by their one-lag covariances, their spatial covariances being identical. For these type of input structure, mean-LP and cov-LP alone (i.e. without reservoir) cannot capture the relevant statistics for classification, so they produce chance level accuracy. The reservoir thus becomes fundamental for this task. As with the spatial structure, we find that performance increases with reservoir size for both readout types. However, the performance for the covariance-based decoders is well above that of mean-based ones, and approximates that of the MLR directly applied to the one-lag covariances of the inputs (98.8 ± 0.5%, dashed light gray line). Thus, the conversion from temporal second-order patterns in the inputs to spatial second-order patterns in the reservoir is more efficient than to spatial first-order patterns. The temporal structure yields different accuracy dependencies on the spectral radius and leak rate, as compared to the spatial structure. First, we note that mean-based readouts perform very poorly for all tested configurations and that both decoders achieve their best performance for non-zero spectral radii. This indicates that the reservoir recurrent dynamics are essential to transform the input lag covariances into output spatial statistics of the reservoir activity, of either first or second order. The optimal reservoir configurations have recurrent connectivity with *ρ*(Ω^res^) ≈ 1, which have also been shown to maximize memory capacity for Gaussian inputs (Jaeger, 2002; Farkaš et al., 2016). In those studies, the reservoir transforms dynamic signals in a way that allows to retrieve past information for a given range of delays. Instead, we here do the converse and transform the temporal structure (lag covariances) into a spatial structure (zero-lag covariances). For a reservoir in a linear or weakly nonlinear regime, indeed it can be shown that optimal representations at both orders are obtained when *ρ*(Ω^res^) → 1 from below (see Section2.1.1 in Methods). Nonetheless, as these temporally structured inputs produce reservoir nonlinear dynamics that are stronger than the ones induced by spatially structured inputs (Fig. 3B), representation degradations seem much more pronounced at the first-order than the second-one, explaining the larger gap in accuracy between mean and covariance based decoders in Fig. 3A.

Second, accuracy increases with leak rate for both readout types. Nonetheless, for covariance readouts in a feedforward reservoir with *ρ*(Ω^res^) = 0, leaky integration is key, as it allows neurons to keep a memory of past inputs in their current state.

We further compare these LP decoders to the MLR decoders, in both mean-based and covariance-based versions as illustrated in Fig. 3C. As before, we focus on the case where a reservoir is involved in the classification pipeline (RES-N, corresponding to the bottom pathway in Fig. 2A), as well as the decoders directly applied to the inputs (NO-RES, see the top pathway in Fig. 2A). The best decoder performance for each case in Fig. 3A across the considered reservoir parameters is displayed in Fig. 3C. It can be seen that covariance-based decoders outperform mean-based ones when the information to extract is in the second-order statistics of the inputs across all models tested, and that the best decoder is the cov-MLR, which is also the readout with highest complexity (number of parameters).

Ultimately, temporal and spatial input structures are efficiently processed by reservoirs, but distinct characteristics give the best performance (especially the radius). A good compromise for both types of inputs and covariance-based readouts is a reservoir without leaky integration and medium-sized spectral radius, namely *α* = 1 and *ρ*(Ω^res^) ≃ 1.

**Figure 4:**
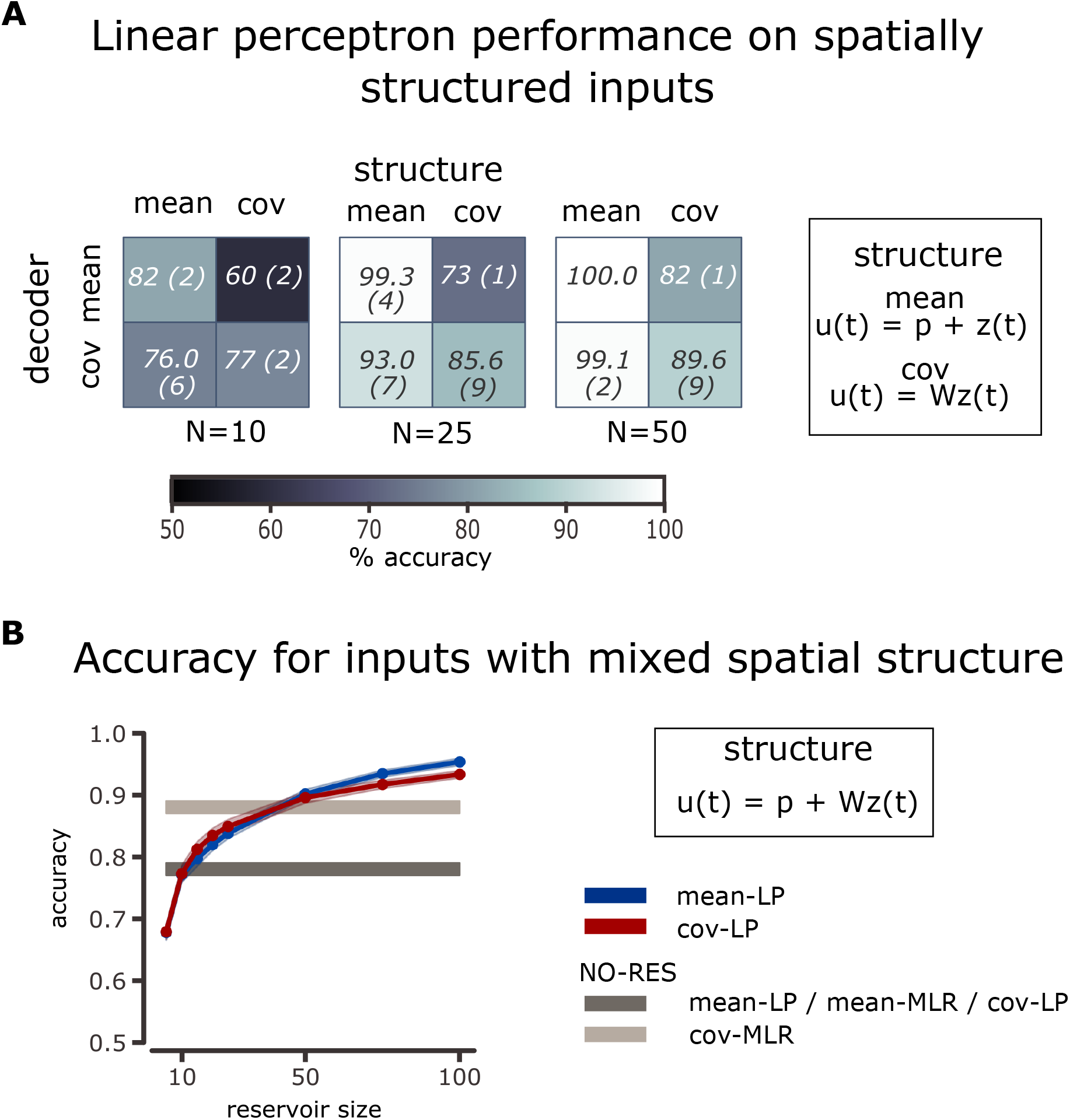
Decoding mixed information for the mean structure and spatial structure using feedforward reservoirs. **A:** Accuracy table for feedforward reservoirs without leaky integration coupled to a linear perceptron when using mean or covariance readouts to classify time series characterized by its mean or its spatial covariance structure. We focus on small reservoirs (*N* = 10, 25, 50), since the mean structure task gives perfect performance for such reservoir size (*N* ≥ 50). Each entry in the table is the mean accuracy (with sem in parentheses) across 10 different simulations. Datasets were designed to match the performances using the perceptron as readout: cov-LP 79(±2)%, 15 patterns per class, mean-LP 81(±2)%, 10 patterns per class. **B:** Classification accuracy as a function of reservoir size for a feedforward reservoir without leaky integration coupled to a linear perceptron as a readout when the categories of input time series differ by both their means and their spatial covariances. As before in Fig. 3A, the performance for the mean-LP is represented in blue, that of the cov-LP in red. Shaded areas represent ±1 sem across 10 different simulations and gray lines indicate the performance of decoders that are directly applied to the inputs (NO-RES). Note that the datasets are generated in such a way that the classification performance of a mean-LP and cov-LP without reservoir is matched, equal to 78 ± 1%, with 10 patterns of each statistic to distinguish per class (see Section 2.3.4 in Methods).

### 3.2 Reservoirs with covariance-based readouts can also extract firstorder statistics

Thus far, we have shown that a consistent processing scheme between inputs and outputs via the reservoir for covariance-based information processing is better than a mixed scheme that maps input covariances to means in the reservoir activity, which are then used by the readouts for decoding. Now, we explore for comparison the case where the inputs have embedded information in their mean activity. When class information is embedded in spatial statistics, we have shown analytically and numerically that a pointwise nonlinearity is key to produce reservoir representations shaped at first and second order, while the recurrent dynamics are less important. Thus, we restrict this investigation to a feedforward reservoir and no leaky integration, namely *ρ*(Ω^res^) = 0 and *α* = 1, as they give the best performance in Fig. 3A. Note that the leak rate is less crucial than the absence of recurrent connectivity here.

We firstly compare the performance of mean-LP and cov-LP decoders when the input information is embedded in two different statistical orders: either means (input dynamics in Section 2.3.1) or spatial covariances (input dynamics in Section 2.3.2). We intentionally work with small reservoirs to avoid the case where the mean classification task becomes trivial, with perfect performance. To fairly compare the two structure schemes, we create the respective datasets such that the performance of a LP classifier acting on the distinctive input statistics (either mean or covariances) is matched up to 1% of performance. Our results in Fig. 4A indicate that, typically, the best strategy is to use for decoding the same order the input is structured at, for small reservoirs. However, when the reservoirs are big enough (*N* ≥ 50), switching to mean-based decoding can be beneficial, as learning is computationally faster.

Then, we consider the situation where the input time series can be categorized by either of their first/second-order statistics, corresponding to the dynamics in Section 2.3.4. In this setting, knowing only one of these statistics is enough to perform the binary classification, and the question is whether a type of decoding can make use of both types of information in a synergistic manner. As before, the inputs are designed to match decoding performances when LP is applied directly to them. We find that both decoders perform equally well across various reservoir sizes (Fig. 4B), with a slight advantage for cov-LP over mean-LP for smaller sizes and conversely for larger sizes.

These results are consistent with the fact that in a feedforward reservoir, if input-to-reservoir weights are 𝒪(*ε*), with |*ε*| *<* 1, then the statistical moments of reservoir neurons will depend on input statistics with different leading orders in *ε* (see Methods 2.1.1).We have shown that reservoir mean activity has 𝒪(*ε*) dependence on input mean activity and 𝒪(*ε*^2^) dependence on input spatial covariances. On the other hand, reservoir spatial covariances display the same 𝒪(*ε*^2^) dependence on both input statistics. Thus, if information is on input means, it is more strongly reflected on reservoir means, and likewise for covariances. However, when information is embedded in both statistical orders, then fixing a representation for the reservoir settles the other one as ‘noise’. Therefore, in a mixed scheme, mean-based representations have a better signal-to-noise ratio than covariance-based ones.

Together, these results show that the application of covariance-based decoding, when combined with a reservoir, goes beyond that of its same order encoding: information about multiple input statistical orders can be effectively mapped to output second-order moments.

### 3.3 Covariance-based information is more efficiently propagated in short time scales

As a last point with the synthetic data, we consider a reservoir with separated input receptor and output feeder units (Section 2.1.3) to study the role of the reservoir topology in the transmission of different statistical orders (Fig. 2B). In this architecture motivated by biology with segregated functions, signals have to propagate from end-to-end of the reservoir to reach the decoder in a short-time window *d* = 20. We compare segregated reservoirs with different sparse connectivities (Fig.2B), as well as the architecture studied until now with full connectivity from inputs and to outputs. Importantly, we design these configurations such that they share the same number of decoder resources, as given by the number of connections from reservoir units to output units (Fig. 3A, middle row, *N* = 50). The leak rate is always set to *α* = 1. Since the density of within reservoir connections does not affect performance in the full connectivity configuration (it is only important that it is non-null, see Supplementary Material Fig. 8), we set the reservoirs to have 0.1 density.

For the task of decoding input spatial covariances, we observe that the performance is improved for mean-decoding (Fig. 5A, left panel, blue) with large spectral radii when the segregated reservoir has symmetric structure. On the other hand, covariance-based decoding is only reaching the not segregated reservoir performance for medium radii (*ρ*(Ω^res^) = 0.9), where the segregation of receptor and feeders impedes that this task be achieved by a feedforward reservoir.

**Figure 5:**
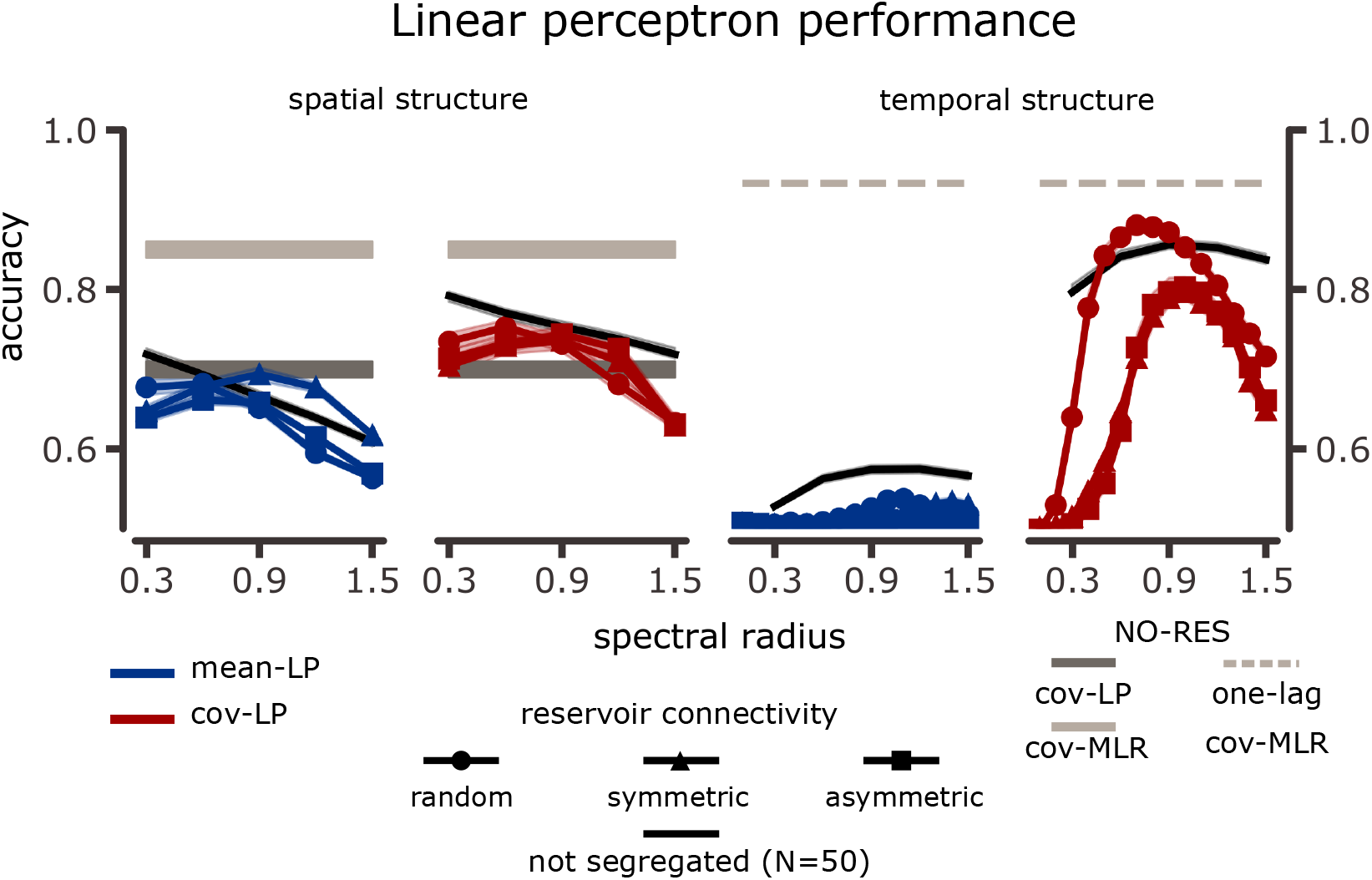
Decoding performance for spatial and temporal structure in segregated reservoirs. Segregated reservoir (Fig 2B) classification performance of spatial structure (left panel) and temporal structure (right panel) embedded in the input time series when the decoder is a linear perceptron. Accuracy is shown as a function of spectral radius and for different reservoir topologies (marker styles). Color coding is similar to Fig. 3A. The black lines show the performance of a non-segregated reservoir of 50 neurons without leaky integration (*α* = 1, same as in Fig. 3A, middle row) for comparison.

When the information to decode is in the temporal covariances (Fig. 5A, right panel), mean-LP performance is degraded. Covariance decoding, on the other hand, is better preserved and even reaches a better performance than a fully connected random topology for *ρ*(Ω^res^) ≈ 0.7. For symmetric and asymmetric connectivities, accuracy is significantly degraded across all spectral radii when compared to the not segregated reservoir, and we also find that the relationship between optimal performance for these topologies and spectral radius changes. This suggests a complex interplay between the input structure and the reservoir topology, which would be interesting to explore in a more analytical manner in future work.

On average, information about input covariances is better recovered by covariance decoders even when the patterns must propagate along a network to reach the output units in a short time window.

### 3.4 Covariance-based schemes work in classification of spoken digits

Finally, we explore the applicability of covariance-based readouts in reservoir computing for the classification of spoken Arabic digits (Hammami and Sellam, 2009; Hammami and Bedda, 2010; Dua and Graff, 2019). The dataset consist of digits 0 − 9, represented by 13 input nodes obtained after some preprocessing mimicking the cochlea as illustrated in Fig. 2B (see Section 2.4 in Methods for further details). Our goal here is not so much to improve the best performance obtained so far on this dataset. Rather we aim to evaluate our covariance-based decoding in a real case study and speech recognition is a common area of application for reservoir computing, given the sequential nature of the task (Verstraeten et al., 2005, 2006, 2007; Skowronski and Harris, 2007; Triefenbach et al., 2010; Alalshekmubarak and Smith, 2014; Zhang et al., 2015; Jin and Li, 2017; Aceituno et al., 2020). Indeed, the classical mean-based decoding reaches a competitive performance of 99.9923% for a reservoir of size bounded by *N* = 1, 000 (Aswolinskiy et al., 2018), and it can be further increased to 99.9945% when using a predictive model space representation instead of the natural reservoir space (Aswolinskiy et al., 2018).

As before, we use the two pipelines for classification, with and without reservoir (same as Fig. 1A) and we vary reservoir properties (size, spectral radius and leak rate) across a grid. For brevity, we only report the test accuracy of the best performing models for each reservoir size and decoder in Fig. 6A, which are obtained for *α* = 0.2 and *ρ*(Ω^res^) = 1.2 (see Supplementary Material Fig. 9 for all results). This is in line with decoding temporal structure (Fig. 3A), rather than spatial structure, suggesting that the temporal structure of the real data is best captured by the reservoir to perform efficient classification. First, we note that when we do not use a reservoir, the best decoders are the nonlinear ones (mean-MLR and cov-MLR), and that both statistical orders contain relevant information to classify the time series. However, the higher accuracy obtained with the cov-MLR indicates that the covariance patterns are potentially easier to extract than the mean patterns. Indeed, the performance of cov-MLR with *N* = 100 is 98.7 ± 0.1%, better than using a mean-based decoder with an echo state network of *N* = 900 neurons (96.91%(Alalshekmubarak and Smith, 2013)), but obtained with 9 times fewer neurons within the reservoir. Furthermore, the performance is at the same level of that obtained with a much more complex network model, as is the long short-term memory network in (Zerari et al., 2019) (98.77%).

**Figure 6:**
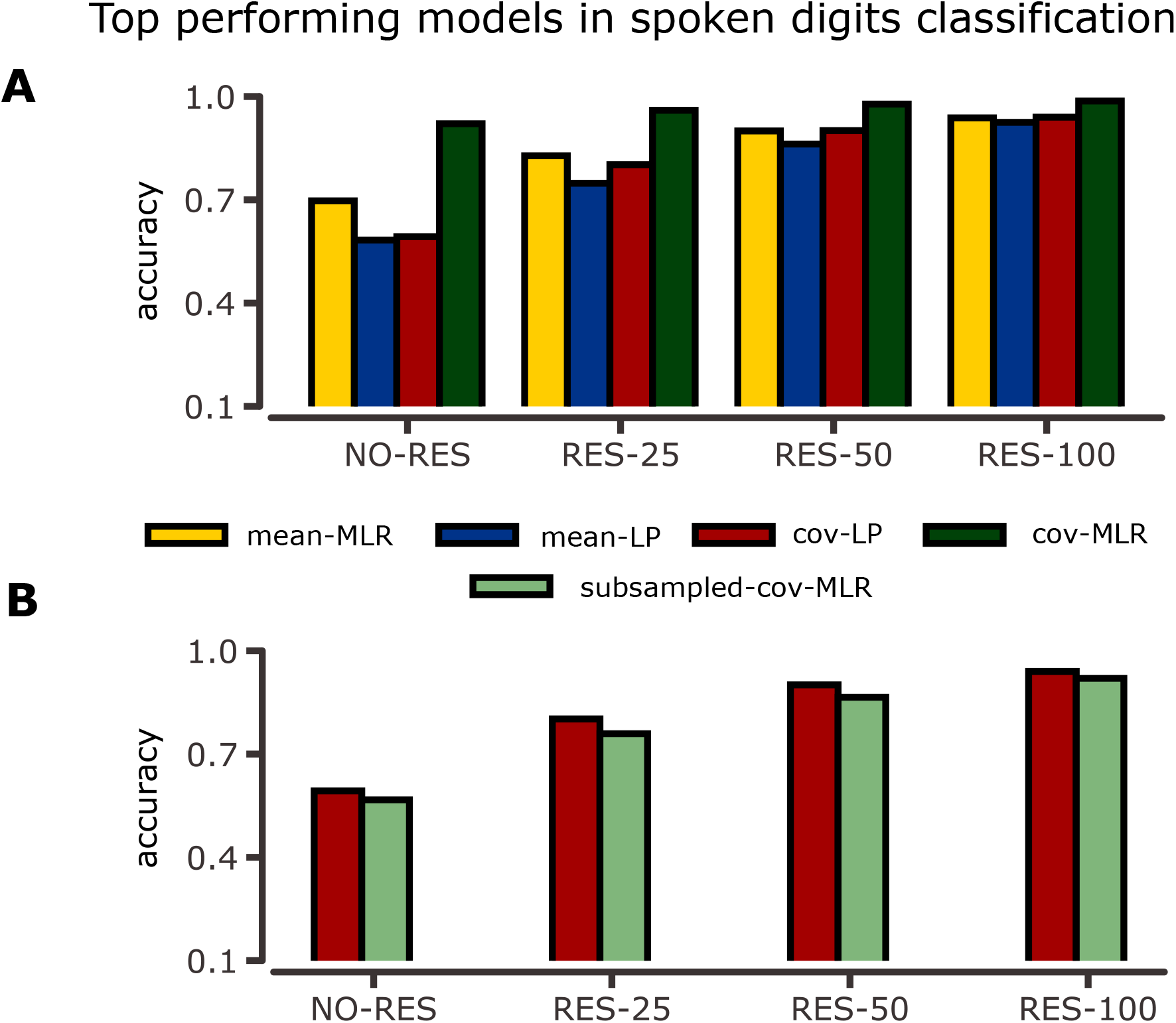
Decoding performance in spoken Arabic digits classification. **A:** Accuracy for the best performing models of each size, distinguished by decoder type. Colors and model label conventions are the same as in Fig. 3B. Results are averaged across 10 different reservoir instantiations. Error bars displaying ±1 sem are also included. **B:** Accuracy for the best performing models with varying reservoir size, when the decoder is covariance-based (cov-LP and cov-MLR) and the number of parameters of each decoder are approximately matched. Error bars displaying ±1 sem are also included.

In addition, the linear decoders (mean-LP and cov-LP) perform well above chance level, which corresponds here to 10% because the dataset is balanced across the 10 possible digits. The use of a reservoir in the pipeline is beneficial to all decoders, and the resulting performance increases with reservoir size. When *N* = 100, the performances of the mean-MLR, mean-LP and cov-LP are closely matched, but they are still below that of the cov-MLR. Our insight is that this gap could be further reduced by increasing the reservoir size.

To further compare the cov-LP and cov-MLR, we subsample the number of neurons in the reservoir, so that its vectorized covariance matrix has dimensionality close to *N*, instead of *N* (*N* + 1)*/*2 (see Section 2.2.2 for details). We find that both models exhibit similar performance under this constrain (see Fig. 6B), which confirms that the cov-LP decoder extracts information about covariances in an optimal manner given its limited number of resources.

All in all, covariance-based decoders can be successfully applied within reservoir computing frameworks to classify spoken digits. The covariance perceptron (cov-LP) applied to reservoir computing offers a good compromise between good performance and limited resources as in a biological context.

## 4 Discussion

In this study, we have explored how a neuronal reservoir can be efficiently paired with covariance-based readouts for the classification tasks of time series. Our goal was to investigate the potential of this new type of decoding as compared to the classical mean-based decoding that has been used with reservoir computing until now, with a two-fold motivation. First, we aimed to characterize how reservoir dynamics can process the statistical structure embedded in the input time series, up to the second order including spatial and temporal structures. Second, we wanted to compare a biologically inspired configuration —a perceptron network mapping time series to time series— with a machine learning configuration —multinomial logistic regression mapping static features to class probabilities. Our results demonstrate the efficiency of covariance-based readouts applied to reservoirs, even in the biological configuration that involves limited resources as implemented by a (linear) covariance perceptron.

We have shown using synthetic data that covariance decoding allows for capturing a broad diversity of input structures after their transformation by the reservoir. To do so, a compromise configuration for our echo state networks, corresponding to *α* = 1 and *ρ*(Ω^res^) = 1, is robust to variations in second-order input structure. These findings are confirmed with the classification of spoken digits from a real dataset, for which we find better performance for covariance decoding: the performance of cov-MLR with *N* = 100 (98.7 ± 0.1%) is better than the mean decoding with *N* = 900 (96.91%) (Alalshekmubarak, 2014; Alalshekmubarak and Smith, 2014). Moreover, the cov-LP maximally extracts information about covariances given its restricted resources. For the real digits dataset, the best radius is similar to the compromise configuration for synthetic data, but the best leak rate is smaller. This is in line with previous work in reservoir computing studies with mean-based readouts, which suggest that small leak rates better suit the intrinsic time scale of the input time series, as speech spectral features vary slowly when compared to the sampling frequency (i.e. the spacing between the windows used to compute MFCC) (Verstraeten and Schrauwen, 2009). In such a case, the usefulness of the reservoir is to transform signals at those slow timescales into zero-lag correlations of the reservoir activity.

In more detail, the collective dynamics of the reservoir, as governed by the interplay between the spectral radius and the leak rate, has a crucial effect on the input-output mapping in terms of statistics. For large radii, small leak rates produce slower dynamics that tend to drive the reservoir towards the linear or weakly non linear regimes, which in turn enhances mean decoding of spatial structure. When inputs are endowed with spatial structure, covariance decoding is less dependent on leak rate, which suggests that reservoir pairwise correlation patterns mostly depend on global dynamic features. On the other hand, mean decoding of temporal structure performs very poorly and is negatively affected by slow unit and global dynamics, as information from the past has to rapidly produce spatial patterns distinguishable by the mean decoder. For covariance decoding when the information source are the one-lag covariances, the global memory mechanism given by the spectral radius and the individual one given by the leak rate operate best when acting alone, as we find that the presence of slow unit dynamics decreases performance when the reservoir is scaled to operate close to the unstable regime (*ρ*(Ω^res^) ≈ 1). The optimal reservoir in this case is one without leaky integration and *ρ*(Ω^res^) ≈ 1, which is balanced between the linear, weakly nonlinear and nonlinear regimes. Thus, driving the system closer to the linear regime by decreasing the leak rate (and effectively reducing the strength of the recurrent connections for a given spectral radius) degrades performance. While most analytical studies in the reservoir computing literature focus on the linear approximation (Jaeger, 2001, 2002; Jaeger et al., 2007; Aceituno et al., 2020), our numerical results suggest that other input-induced dynamical regimes should be further examined theoretically (Manjunath and Jaeger, 2013; Verzelli et al., 2020).

Early studies in echo state networks failed to report performance improvements in time series prediction when using small-world or scale-free topologies (Liebald, 2004; Rad, 2008). However, it was later shown that connectivity plays a role when the reservoir network displays cortex-like topological properties (Song and Feng, 2010; Kawai et al., 2019), a finding that had been observed in reservoirs of spiking neurons earlier (Haeusler and Maass, 2006). In those studies as replicated in ours, nodes receiving inputs differ from those feeding the readouts, thereby mimicking the separation of sensory areas from motor areas. This scheme is most efficient to transform input temporal covariances into output spatial covariances when the reservoir connectivity is random; note that in this case the optimal spectral radius is smaller than for the non-segregated reservoirs (*ρ*(Ω^res^) = 0.7 instead of 1). This could be due to the overall bigger reservoir size that provides a faster mixing of the inputs (*N* = 500). Surprisingly, symmetric connectivity enhances mean decoding of spatial structure. Overall, there is a nontrivial interaction between reservoir topology and input structure which should be further investigated. We note that future work could explore more elaborate topologies, like small-world, scale-free, clustered (Weidel et al., 2020) or even real connectome patterns (Suárez et al., 2020; Damicelli et al., 2021), beyond the simple connectivities studied here. Furthermore, we constrained our study to random input projections, which take part in shaping the input representations that arise in the reservoir (as hinted in Section 2.1.1). Along this line, it has been shown that unsupervised plasticity at the input-to-reservoir layer improves performance in pattern recognition tasks with mean-based decoding (Weidel et al., 2020), so it is natural to question whether this effect is also observed, or even further enhanced, for covariance-based readouts.

Last, we stress that the use of the reservoir here offers several advantages with respect to the linear network studied in (Gilson et al., 2020). First, it avoids the training of recurrent connections in the readout layer to classify temporal covariance structures, by retaining past information in its own activity. Thus, learning is computationally cheaper, as there is no need to numerically solve Lyapunov equations, which is the case in the recurrent covariance perceptron (Gilson et al., 2020). Second, the cross-talk among statistical orders induced by the reservoir allows the covariance perceptron to capture a broader variety of input statistics, in particular when the input information is embedded in the first statistical order. Meanwhile, the neuronal system remains biologically plausible and takes advantages of the interplay between nonlinearities and recurrent connectivity (Maass et al., 2002; Enel et al., 2016). Although we have not examined the influence of the nonlinearity used in the reservoir, we expect a variety of biologically inspired functions to lead to efficient computations, provided they keep the recurrent dynamics under control for medium radii (i.e. bounded activity). Our work may also bring a novel perspective in training recurrent networks with feedback in the line of the FORCE algorithm, where the focus is on generating patterns (time series) that consist of trajectories (Sussillo and Abbott, 2009; Miconi, 2017; Klos et al., 2020). A possible extension is to generate time series with desired covariance-based patterns instead.

Another extension in the direction of biological realism is to transpose the scheme to spiking neurons (Maass et al., 2002). Following from our results when our analogue reservoir is close to a linear regime, we expect our covariance-based decoding framework to give interesting perspectives in terms of operations on covariances for spiking networks in similar linear regimes. However, the specific nonlinearities involved in spiking neurons should have significant effects on the input-output mapping and require a thorough study. Such covariance-based learning for spiking neurons would be an intermediate between learning spike rate patterns and precise spike trains, as with the ‘tempotron’ (Gütig and Sompolinsky, 2006) or ReSuMe (Ponulak and Kasinśki, 2010).

An important limitation of our study has been on the size of the reservoirs we were able to implement, as the perceptron learning procedure for covariances becomes numerically unstable as the number of parameters to tune increases. Current work on overcoming this and studying the scaling behavior of covariance-based reservoirs is focused on using techniques such as gradient-clipping (Pascanu et al., 2013). Nonetheless, this approach makes the learning procedure slow as the number of optimization steps needed to find a good solution increases. Another issue to also address in the future is how to efficiently regularize covariance-based decoders, since going for larger reservoirs might lead to overfitting. These considerations underlie a cost-benefit trade-off between covariance and mean-based representations. While covariances offer higher-dimensional representational spaces than means for the same number of resources, learning is still computationally cheaper and more stable for mean-based representations (Gilson et al., 2020; Dahmen et al., 2020).

## Acknowledgements

S.L is supported by a FI fellowship from the Agència de Gestió d’Ajuts Universitaris i de Recerca (AGAUR, 2021 FI-B2 00121). R.M.-B is supported by the Howard Hughes Medical Institute (HHMI, ref 55008742), MINECO (Spain; BFU2017-85936-P) and ICREA Academia (2016). M.G acknowledges funding from the German Excellence Strategy of the Federal Government and the Länder (G:(DE-82)EXS-PF-JARA-SDS005) and the European Union’s Horizon 2020 research and innovation programme under grant agreement No. 785907 (Human Brain Project SGA2).

## Code availability

Python code samples to generate synthetic datasets and reservoirs, training methods, as well as some pretrained models can be found at http://www.github.com/slawrie/covariance-reservoir.

## A Supplementary Material

### A.1 Reservoirs propagate diverse input statistics

Let *u*(*t*) ∈ ℝ^*M*^ be a multivariate time series fed to an echo state network, with *u*_1_(*t*) a bias unit. The update equations for an arbitrary neuron *x*_*i*_(*t*) inside a reservoir with *N* units are given by:

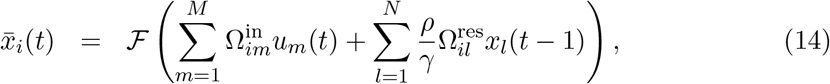

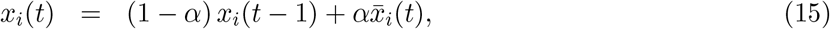

where the function ℱ = tanh has a sigmoidal profile, Ω^in^ ∈ ℝ^N×M^ is the connection matrix from the input time series to the reservoir units and Ω^res^ ∈ ℝ^N×N^ is the weight matrix of recurrent connections within the reservoir. Factor 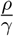 allows to control the spectral radius of the effective matrix of recurrent connections 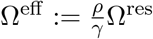, where *γ* is the spectral radius of Ω^res^.

In a fully leaky reservoir (*α* = 1) and assuming a left-infinite sequence of inputs is presented to the reservoir, then the state at time *t* of a neuron can be written as

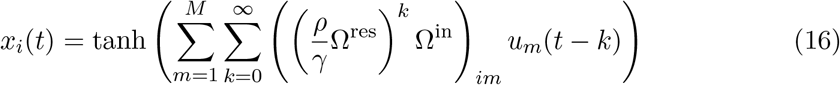

If *α* = 1 and neurons are in a weakly nonlinear activation state, we can approximate Eq. 16 by a Taylor series truncated at the third order:

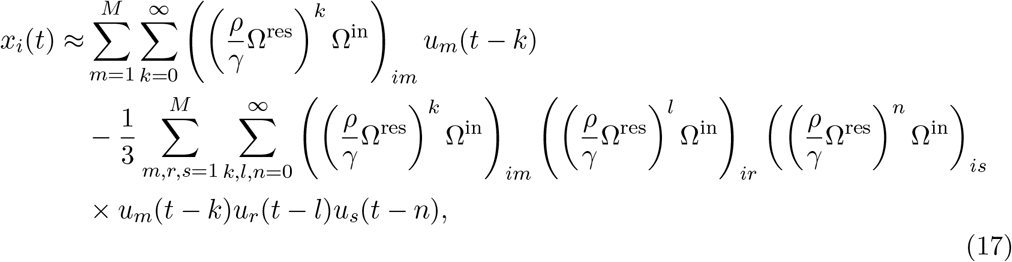

where we have collapsed the sums 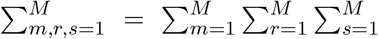 and 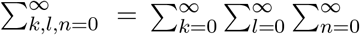 for ease of notation and have used that tanh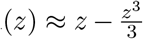 when *z* ≈ 0.

In the following, we will assume:

- Ω^in^ elements are independently sampled from the uniform distribution over [−*ε/M, ε/M*]
- Ω^res^ elements are independent and identically distributed random variables with zero mean and variance 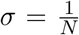, and thus in the limit *N* → ∞, *γ* → 1 (Ginibre, 1965).

We want to observe how reservoir first and second-order statistics are influenced by input statistics. Therefore, note that:

- In magnitude, elements in 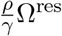 follow 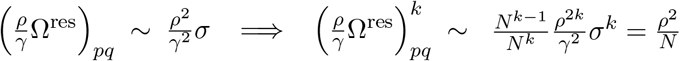
- Elements in Ω^in^ follow 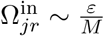
- 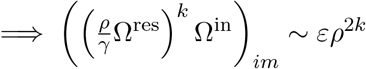

Thus, we get that:

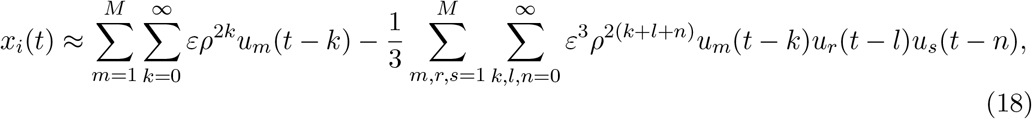

Thus, if we compute the mean ⟨*x*_*i*_(*t*)⟩, we obtain:

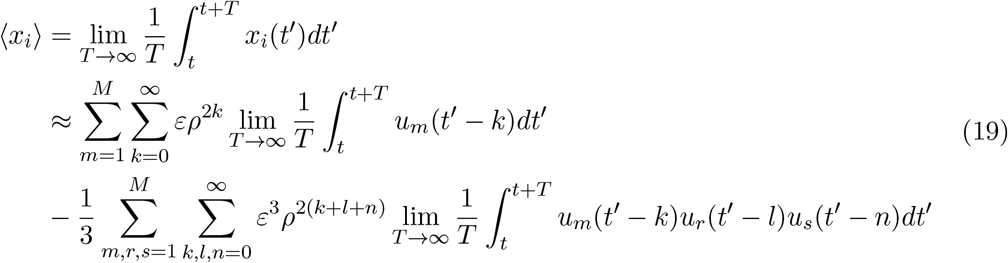

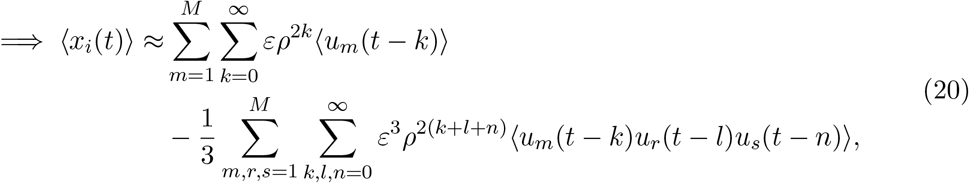

If we assume that the input time series are stationary, then their statistical moments do not change when time shifted, i.e. ⟨*u*_*m*_(*t* − *k*)⟩ = ⟨*u*_*m*_(*t*)⟩ and ⟨*u*_*m*_(*t* − *k*)*u*_*r*_(*t* − *k*)*u*_*s*_(*t* − *k*)⟩ = ⟨*u*_*m*_(*t*)*u*_*r*_(*t*)*u*_*s*_(*t*)⟩ (they are time invariant). Thus, we can compute the infinite sums (which are convergent geometrical series if *ρ <* 1), and get the dependence of ⟨*x*_*i*_(*t*)⟩ in terms of the first order and second (zero and one lag) covariances of the inputs:

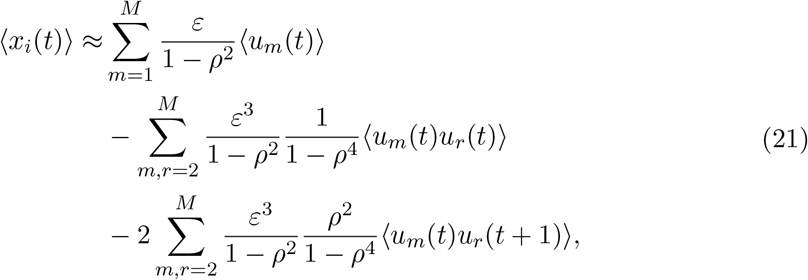

Following the same procedure, we can compute ⟨*x*_*i*_(*t*)*x*_*j*_(*t*)⟩ using the linear approximation to the state of each neuron. Thus, we get:

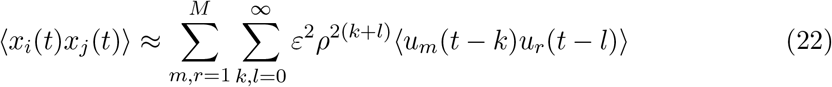

And in terms of first and second order moments (zero and one-lag), we find:

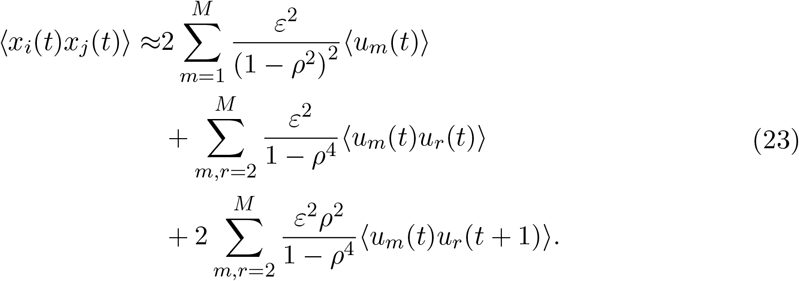

Thus, we observe that *ρ* plays a key role in how input statistics are reflected in reservoir statistics.

### A.2 Section 3.2 results when using MLR decoders as benchmarks

**Figure 7:**
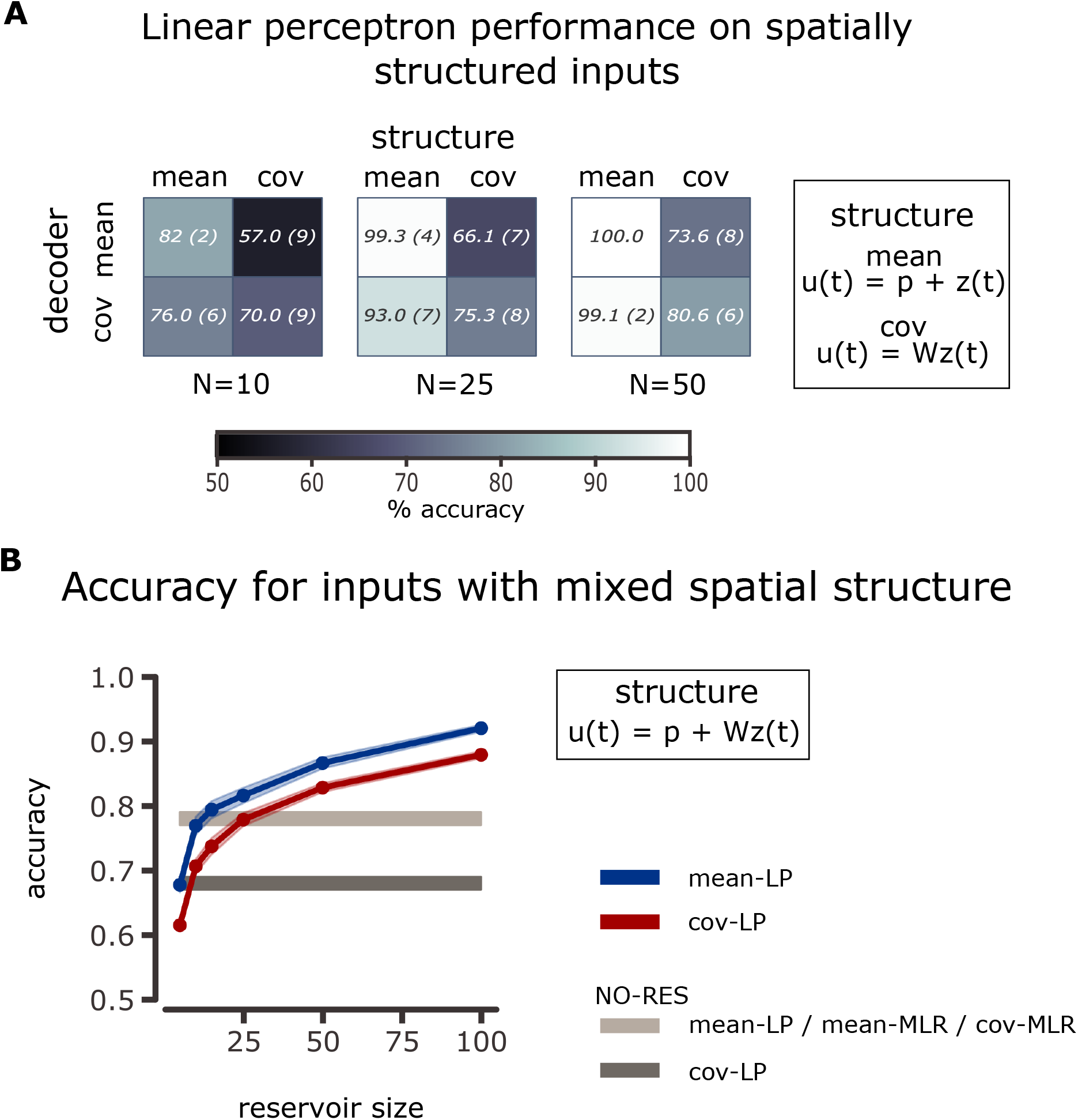
Decoding mixed information for the mean structure and spatial structure using feedforward reservoirs. **A:** Accuracy table for feedforward reservoirs without leaky integration coupled to a linear perceptron when using mean or covariance readouts to classify time series characterized by its mean or its spatial covariance structure. Same conventions as Fig 4. Datasets were designed to match the performances using the MLR as readout: cov-MLR 85(±1)%, 30 patterns per class, mean-MLR 83(±2)%, 10 patterns per class. **B:** Classification accuracy as a function of reservoir size for a feedforward reservoir without leaky integration coupled to a linear perceptron as a readout when the categories of input time series differ by both their means and their spatial covariances. Same conventions as in Fig 4. Note that the datasets are generated in such a way that the classification performance of a mean-MLR and cov-MLR without reservoir is matched, equal to 78±1%, with 10 patterns of mean vectors and 20 patterns of covariances to distinguish per class.

In Section 3.2, we compare the performance of cov-LP and mean-LP decoding when the statistical information is embedded in the first or second-order moments of the inputs (Fig 4a) or in both (Fig 4b). To be able to compare performance across different datasets, we chose the number of patterns to distinguish per class such that mean-LP and cov-LP decoders acting directly on the inputs obtained matching performances. Using instead MLR decoders to match datasets, we can produce the same figure. Note that in this case, for Fig 7a, we have 30 covariance patterns per class (as in Fig 3a, left pannel) instead of 20. For Fig 7b we have 20 covariance patterns per class, instead of the 10 in Fig 4b. There is now a difference between the number of patterns to distinguish in each space, given mainly because cov-MLR decoders have a higher model complexity. Thus, to generate the dataset each mean pattern is randomly paired to 2 covariance patterns, and then the set is randomly assigned to a class. We observe, nonetheless, that the results of Fig 4 are mostly preserved. The difference relies in that in the mixed scenario, cov-LP decoders now lag behind mean-LP across all sizes tested. We venture that this difference is due to two sources. One is that when using mean decoders, each noisy pattern is observed twice when compared with each covariance pattern when using covariance decoders. The other is that resources are low for small reservoirs when the input covariance space needs to be covered. This could be overcomed with increasing reservoir size. We observe such decrease in the performance gap between decoders up to 25 neurons. Afterwards, the decreasing stops. This might be due to numerical instabilities when trying to optimize covariance decoders for large number of weights. Overall, Fig 7 suggests that the first-order information overrides the second-order one.

**Figure 8:**
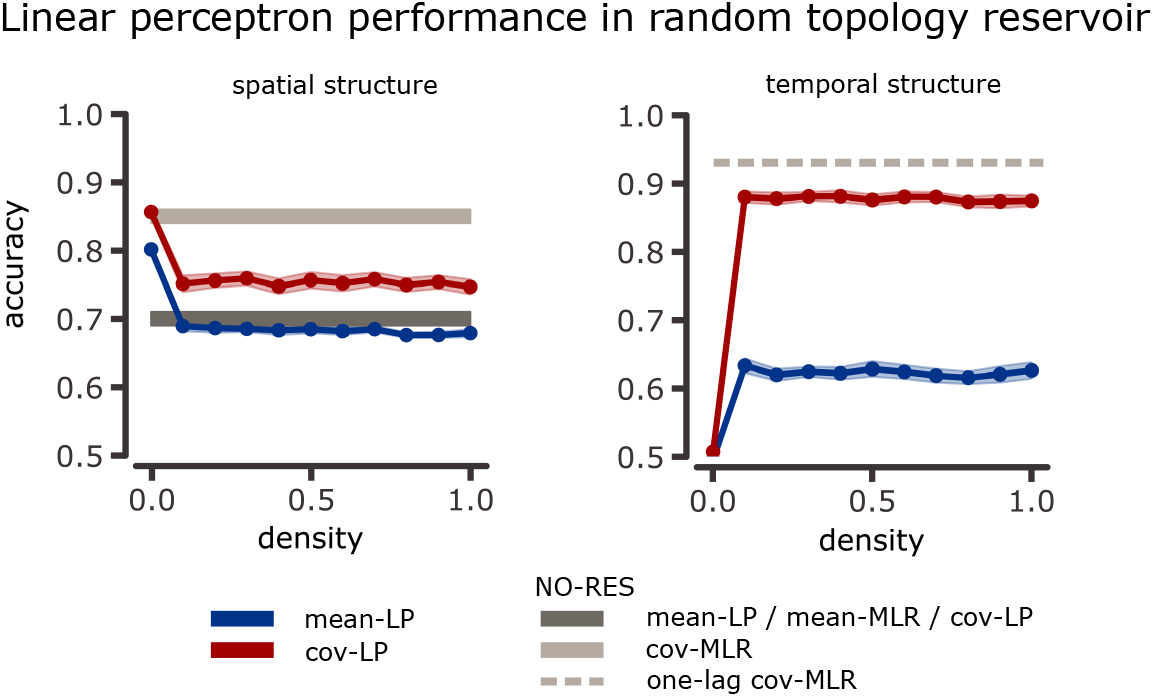
Linear perceptron classification performance in synthetic time series with second-order structure. Similar color conventions as in Fig 3A. Reservoir parameters are *N* = 100, *α* = 1 and *ρ*(Ω^res^) = 1.2.

**Figure 9:**
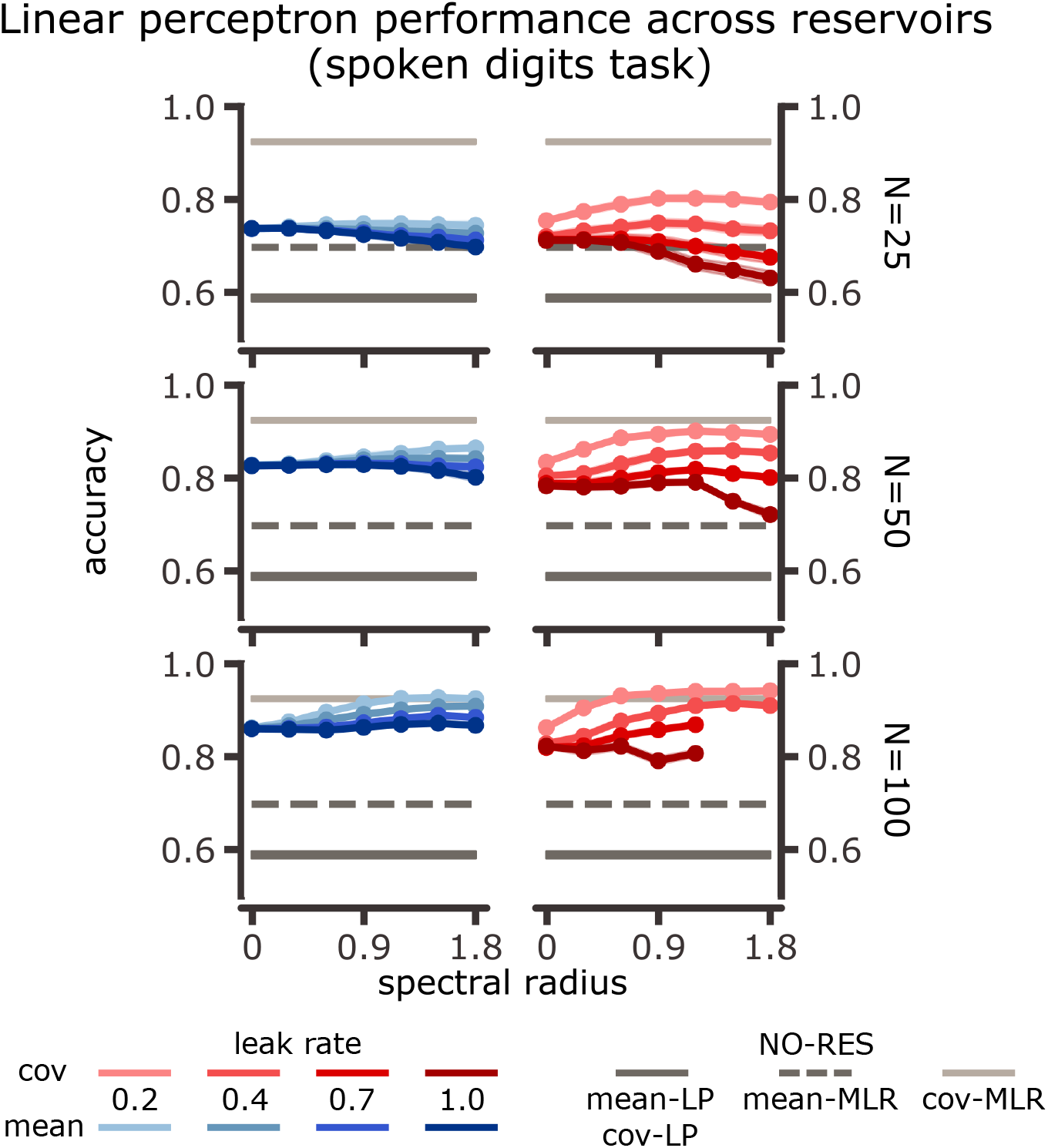
Linear perceptron classification performance for spoken digits across reservoir configurations. Similar color conventions as in Fig 5A. Missing points for *N* = 100, large leaks and spectral radii are due to numerical instabilities during covariance learning.

**Figure 10:**
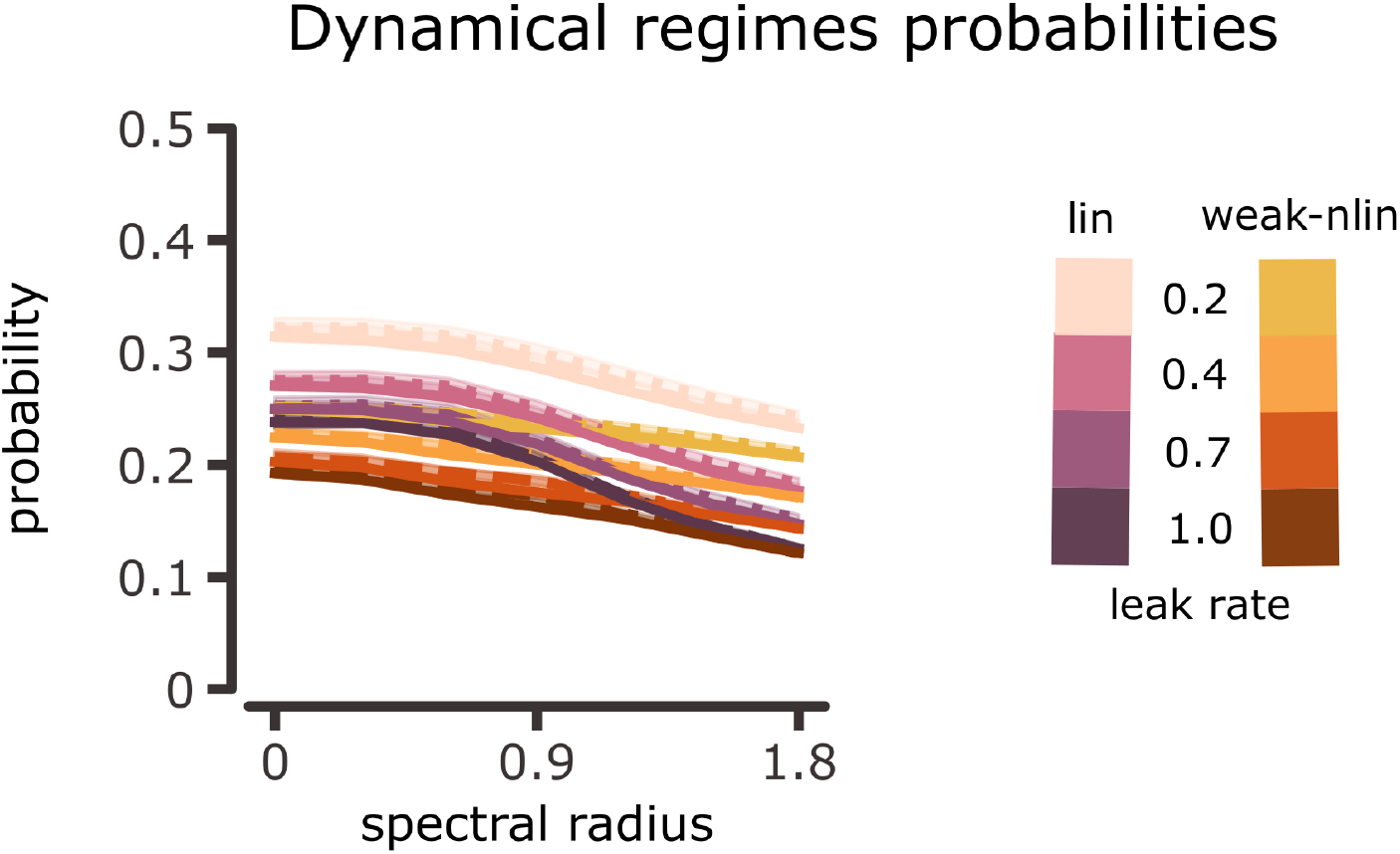
Dynamical regime probabilities for the spoken digits task. Probability of finding a reservoir neuron in the linear or weakly nonlinear activation regimes versus spectral radius, for different reservoir sizes (overlapping dotted, dashed and solid lines for *N* = 25, *N* = 50 and *N* = 100 respectively) and leak rates (shades of gray).

